# Using deep learning to predict internalizing problems from brain structure in youth

**DOI:** 10.1101/2024.11.28.625869

**Authors:** Marlee M. Vandewouw, Bilal Syed, Noah Barnett, Alfredo Arias, Elizabeth Kelley, Jessica Jones, Muhammad Ayub, Alana Iaboni, Paul D. Arnold, Jennifer Crosbie, Russell J Schachar, Margot J Taylor, Jason P. Lerch, Evdokia Anagnostou, Azadeh Kushki

## Abstract

Internalizing problems (e.g., anxiety and depression) are associated with a wide range of adverse outcomes. While some predictors of internalizing problems are known (e.g., their frequent co-occurrence with neurodevelopmental (ND) conditions), the biological markers of internalizing problems are not well understood. Here, we used deep learning, a powerful tool for identifying complex and multi-dimensional brain-behaviour relationships, to predict cross-sectional and worsening longitudinal trajectories of internalizing problems. Data were extracted from four large-scale datasets: the Adolescent Brain Cognitive Development study, the Healthy Brain Network, the Human Connectome Project Development study, and the Province of Ontario Neurodevelopmental network. We developed deep learning models that used measures of brain structure (thickness, surface area, and volume) to (a) predict clinically significant internalizing problems cross-sectionally (*N*=14,523); and (b) predict subsequent worsening trajectories (using the reliable change index) of internalizing problems (*N*=10,540) longitudinally. A stratified cross-validation scheme was used to tune, train, and test the models, which were evaluated using the area under the receiving operating characteristic curve (AUC). The cross-sectional model performed well across the sample, reaching an AUC of 0.80[95% CI: 0.71,0.88]. For the longitudinal model, while performance was sub-optimal for predicting worsening trajectories in a sample of the general population (AUC=0.66[0.65,0.67]), good performance was achieved in a small, external test set of primarily ND conditions (AUC=0.80[0.78,0.81]), as well as across all ND conditions (AUC=0.73[0.70,0.76]). Deep learning with features of brain structure is a promising avenue for biomarkers of internalizing problems, particularly for individuals who have a higher likelihood of experiencing difficulties.

## Introduction

Mental health problems in children and youth are widespread, with 11% estimated to have a mental health disorder^1^. Internalizing problems, specifically, which include feelings of anxiety, depression, and social withdrawal, are associated with profoundly negative outcomes, including poorer quality of life and worse long-term social and economic outcomes^2,3^. This highlights the need for biological markers that help us understand who is most at risk for developing problems in this domain, taking a step towards improving our ability to deliver proactive care that can effectively minimize adverse outcomes. However, these markers have thus far remained elusive.

Neurobiological markers are a promising approach to bridging the gap between wide ranges of genetic and environmental variation and subsequent manifestations of internalizing problems. Measures of brain structure, such as cortical thickness, cortical surface area, and cortical and subcortical volume, are particularly appealing in this context due to their high test-retest reliability^4^. Recent large-scale studies in community samples have shown that internalizing symptoms in youth are associated with altered cortical thickness and volume in frontal, limbic, and temporal regions^5–7^, although some studies have reported these alterations are not specific to internalizing symptoms^8,9^. While these structural markers reveal associations with internalizing symptoms, they offer limited insight into individual-level risk; neuroimaging based-predictive models offer a promising avenue for translating these patterns into tools that can inform personalized mental health care^10^. However, investigations into whether measures of brain structure can predict internalizing problems have been mixed^11–16^. For example, some have reported that children with lower cortical thickness experienced steeper declines in internalizing symptoms over time^11^, while others have reported no predictive effects^13–15^.

Thus far, this work has largely relied on methodological approaches that may be insufficient for characterizing the multi-dimensional and complex brain-behaviour associations related to mental health. For example, regression has typically been used, relying on manual feature selection, restricting brain-behaviour relationships to being linear, and modeling each brain region independently. This calls for complementary strategies which add depth to the existing work, advancing our understanding of the brain-behaviour associations of internalizing problems. Artificial intelligence, in particular deep learning, is increasingly recognized as a valuable tool for predicting mental health outcomes^17^. However, work in this space has primarily focused on predicting diagnostic categories^18^, which overlooks individualized neurobiological variation that is critical for personalized mental health care. While some studies have begun using this approach to predict mental health symptoms from neurobiology^19,20^, it has yet to be applied to predict internalizing problems both cross-sectionally and longitudinally using measures of brain structure.

Here, we used deep learning to predict internalizing problems in youth from neurobiological measures. We drew from four multi-national, independently collected datasets of children and youth from Canada and the United States. We first used cross-sectional measures of brain structure to predict the presence of clinically significant internalizing problems (*N*=14,523), to provide insight into the underlying brain features beyond what can be revealed by traditional approaches. Next, we used longitudinal data (*N*=10,540) to examine whether brain structure can predict subsequent worsening trajectories of internalizing problems. The cohorts include both neurotypical (NT) children and youth and those with neurodevelopmental (ND) conditions. Given the higher prevalence of internalizing problems in the ND compared to NT population^21^, we also provide measures of model performance stratified by the presence or absence of any ND diagnosis alongside overall performance.

## Methods

### Participants

Data from children and adolescents were extracted from four independent datasets: the Adolescent Brain Cognitive Development (ABCD^22^) study (Release 5.1), the Healthy Brain Network (HBN^23^; Release 10), the Human Connectome Project Development (HCP-D^24^) study (Release 2.0), and the Province of Ontario Neurodevelopmental (POND) network (exported January 2024). The ABCD and POND datasets contain longitudinal data, while HBN and HCP-D provide only cross-sectional data. The recruitment strategies, assent/consent procedures, institutional review board approvals, and data usage agreements for the datasets are provided in the **Supplementary material**). Participants were eligible for inclusion based on the availability of T1-weighted structural magnetic resonance images (MRIs) and the Child Behaviour Checklist (CBCL^25^), resulting in at least one datapoint from 14,950 participants (ABCD: 11,796, HBN: 1,959, HCP-D: 498, POND: 697). Flowcharts of the study samples are presented in **Supplemental Figure 1**. The Holland Bloorview Kids Rehabilitation Hospital’s research ethics board approved the current study (#1854).

We acknowledge that individuals have different language preferences for referring to neurodevelopmental disorders/conditions and we honor such perspectives. Given that in this study we defined group membership according to the diagnostic criteria in the DSM-5, we have used this nomenclature in the manuscript, and in partnership with our family and youth advisories (represented by NB and AA).

### Data

Biological sex, race/ethnicity, annual household income, and highest household education were collected as part of all four study protocols (see ***Sociodemographic assessments*** in the **Supplementary material**).

Internalizing problems were measured using the CBCL^25^, a caregiver-reported assessment of problem behaviour in children and adolescents. The assessment derives syndromic scales corresponding to behavioural problems, including anxious/depressed, withdrawn/depressed, and somatic complaints, which are used to calculate the higher-order construct of internalizing problems. Raw scores were converted to age- and sex-normed standardized *T*-scores, with higher scores indicating greater problems. We used binarized outcomes rather than continuous symptoms scores to reduce the influence of measurement error and focus on clinically meaningful thresholds. While continuous scores can vary within a range that may not reflect true differences in severity, binary outcomes based on validated cut-offs help mitigate this issue. Additionally, binary models yield probabilistic outputs, allowing flexible interpretation and alignment with prior work in this area. For the cross-sectional analysis, the provided threshold (*T*-score ≥64) was used to categorize individuals as having clinically significant internalizing problems. For the longitudinal analysis, the reliable change index (RCI^26^) was used to categorize individuals as having worsening (RCI>1.96) or non-worsening (RCI≤1.96) trajectories of internalizing problems (see the ***Reliable change index*** section in the Supplementary material).

T1-weighted images were acquired as part of all four study protocols (**Imaging protocols** in the **Supplementary material**). The FreeSurfer image analysis suite^27^ was used to perform cortical reconstruction and volume segmentation of the structural MRIs (see the ***Imaging preprocessing*** section of the **Supplementary material**), which provided measures of cortical thickness, surface area, and volume according to the Desikan-Killiany parcellation^28^, alongside volumes of subcortical structures, ventricles, white matter, brainstem, cerebellum, and whole-brain. Processed FreeSurfer outputs were provided by ABCD and HCP-D, while the pipeline was run in-house for POND and HBN. Datapoints with poor data quality were excluded from all subsequent analyses (see **Supplementary material**, section ***Quality control*)**.

After quality control, for the cross-sectional analysis, one datapoint per participant was selected, resulting in a sample size of 14,523 children and youth (ABCD: 11,632, HBN: 1,785, HCP-D: 498, POND: 608). For the longitudinal analysis, partial data were available for a third timepoint in the ABCD study, which was leveraged to maximize the total number of included datapoints, resulting in 10,532 datapoints (ABCD: 10,491, POND: 49) from 8,103 participants (ABCD: 8,062, POND: 49) were included. See the ***Datapoint selection*** section in the **Supplementary material** for further information on how datapoints were selected for both analyses.

### Deep learning models

The primary goal of the cross-sectional deep learning model was to establish the features of brain structure that contribute to clinically significant internalizing problems. To do so, we built a binary classification model, a type of model that predicts one of two possible outcomes (here, clinically significant versus non-significant internalizing problems problems), using features of brain structure as input. We then examined the contribution of each feature to the model’s prediction. The model was tuned, trained, and tested using data aggregated across all four datasets. The model architecture was a multilayer perceptron, consisting of an input layer, blocks of hidden layers, and an output layer (see ***Deep learning model architecture*** in the **Supplementary material** for further details). Input features included the measures of brain structure (*N*=241) along with age and sex, given their association with internalizing problems^29^. Note that we did not include ND diagnosis as an input feature, to allow neurobiological patterns to emerge without imposing diagnostic boundaries, consistent with findings of transdiagnostic, data-driven studies that have highlighted a misalignment between diagnostic labels and neurobiological features frameworks ^30–34^. We also ran the model only including age and sex as predictors, to evaluate the added predictive power of neuroanatomy. All features were *z*-scored prior to building the model. A nested 5-fold stratified cross-validation scheme was used to tune, train, and test the model (**Supplemental Figure 3**). For each iteration, the dataset was split into a training set (80%), which was used for hyperparameter tuning and model training, and 20% was withheld for testing purposes. Prediction scores (probabilities of the positive class) and feature importances (Shapley Additive exPlanations^35^ (SHAP) values) were obtained for the test set. To ensure generalizability, this was performed across five folds.

The primary goal of the longitudinal deep learning model was to predict whether an individual would experience worsening internalizing problems from baseline measures of brain structure. The model architecture and tuning, training, and testing procedure was identical to the cross-sectional model. Predictors included baseline age, baseline sex, baseline internalizing problems, and between-timepoint age differences as features alongside the baseline measures of brain structure. Given the sample size imbalance between ABCD and POND, tuning, training, and testing were performed using the ABCD dataset, holding out POND as an external testing set.

### Performance evaluation

The primary measure used to evaluate model performance was the area under the receiving operating characteristic (ROC) curve (AUC). The secondary measure was accuracy, computed as the prediction score cut-off that maximized the average recall of both classes. Outcome measures were computed across all samples, as well as stratified by the presence or absence of any ND diagnosis. Feature importances (SHAP values) were used to measure each feature’s contribution to the model’s prediction.

### Statistical analysis

AUCs and accuracies are reported across folds using the mean and 95% confidence intervals (CIs). Bias with respect to sociodemographic factors (sex, age, race/ethnicity, annual household income, and highest household education) was assessed by examining differences in AUC. Overall feature importances were computed across folds by taking the mean of the magnitudes of the SHAP values across all datapoints.

## Results

### Samples

Data from 14,523 children and youth between 5 – 21 years of age were included for the cross-sectional analysis; participant demographics are presented in **Table 1**. Across all four datasets, 1,967 [14%] participants had clinically significant internalizing problems, and internalizing problems were more prevalent in individuals with an ND diagnosis (31%) compared to those without (8%).

**Table 1:** Participant demographics for the cross-sectional analysis.

|  |  | N [%]<br>ABCD | HBN | HCP-D | POND |
| --- | --- | --- | --- | --- | --- |
| **N** |  | 11,632 | 1,785 | 608 | 498 |
| **ND diagnosis<sup>1</sup>** |  | 1,650 [14] | 1,310 [74] | 0 [0] | 408 [67] |
| **Internalizing problems<sup>2</sup>** | Total sample | 1,129 [10] | 560 [31] | 30 [6] | 248 [41] |
|  | No ND diagnosis | 691 [7] | 130 [28] | 30 [6] | 54 [27] |
|  | ND diagnosis | 435 [26] | 428 [33] | – | 194 [48] |
| **Age range (years)** |  | 8 – 15 | 5 – 21 | 6 – 17 | 5 – 19 |
| **Median age (years) [IQR]** |  | 10.3 [9.5, 11.4] | 10.1 [8.2, 13.1] | 13.0 [10.2, 14.9] | 11.7 [9.7, 14.4] |
| **Sex** | Male | 6,051 [52] | 1,143 [64] | 229 [46] | 430 [71] |
|  | Female | 5,581 [48] | 642 [36] | 269 [54] | 178 [29] |
| **Race and ethnicity<sup>3</sup>** | Hispanic/Latino | 2,368 [20] | 301 [17] | 40 [8] | 23 [4] |
|  | Non-Hispanic Asian | 245 [2] | 50 [3] | 23 [5] | 29 [5] |
|  | Non-Hispanic Black | 1,719 [15] | 191 [11] | 33 [7] | 8 [1] |
|  | Non-Hispanic White | 6,077 [52] | 750 [42] | 308 [62] | 293 [48] |
|  | Multi-Racial/Other | 1,222 [11] | 275 [15] | 89 [18] | 86 [14] |
|  | Unknown/Not reported | 1 [0] | 218 [12] | 5 [1] | 169 [28] |
| **Annual household income<sup>4</sup>** | <\$50,000 | 3,002 [26] | 287 [16] | 61 [12] | 81 [13] |
| | \$50,000-\$99,999 | 2,924 [25] | 322 [18] | 105 [21] | 83 [14] |
| | ≥\$100,000 | 4,705 [40] | 719 [40] | 298 [60] | 197 [32] |
|  | Unknown/Not reported | 1,001 [9] | 457 [26] | 34 [7] | 247 [41] |
| **Highest household education<sup>5</sup>** | Below high school | 429 [4] | 45 [3] | 6 [1] | 2 [0] |
|  | High school or equivalent | 1,930 [17] | 285 [16] | 49 [10] | 41 [7] |
|  | Undergraduate | 3,213 [28] | 570 [32] | 182 [37] | 236 [39] |
|  | Graduate | 2,849 [24] | 854 [48] | 261 [52] | 121 [20] |
|  | Unknown/Not reported | 3,211 [28] | 31 [2] | 0 [0] | 208 [34] |

ABCD: Adolescent Brain Cognitive Development study; HBN: Healthy Brain Network; HCP-D: Human Connectome Project Development study; POND: Province of Ontario Neurodevelopmental network; IQR: interquartile range; <sup>1</sup>According to the DSM-5

For the longitudinal analysis, 10,540 datapoints were included (**Error! Not a valid bookmark self-reference.**). Participants were aged 10.3 [9.6, 10.9] years at baseline (range: 7-15 years), and 12.4 [11.6, 13.2] at follow-up (range: 10-19 years), and the distribution of age at both timepoints is visualized in **Supplemental Figure 2**. Of these participants, 2,029 [19%] of which had worsening changes in internalizing problems between the baseline and follow-up timepoints. The prevalence rates of worsening trajectories in individuals with (17%) and without (20%) an ND were similar.

**Table 2:**
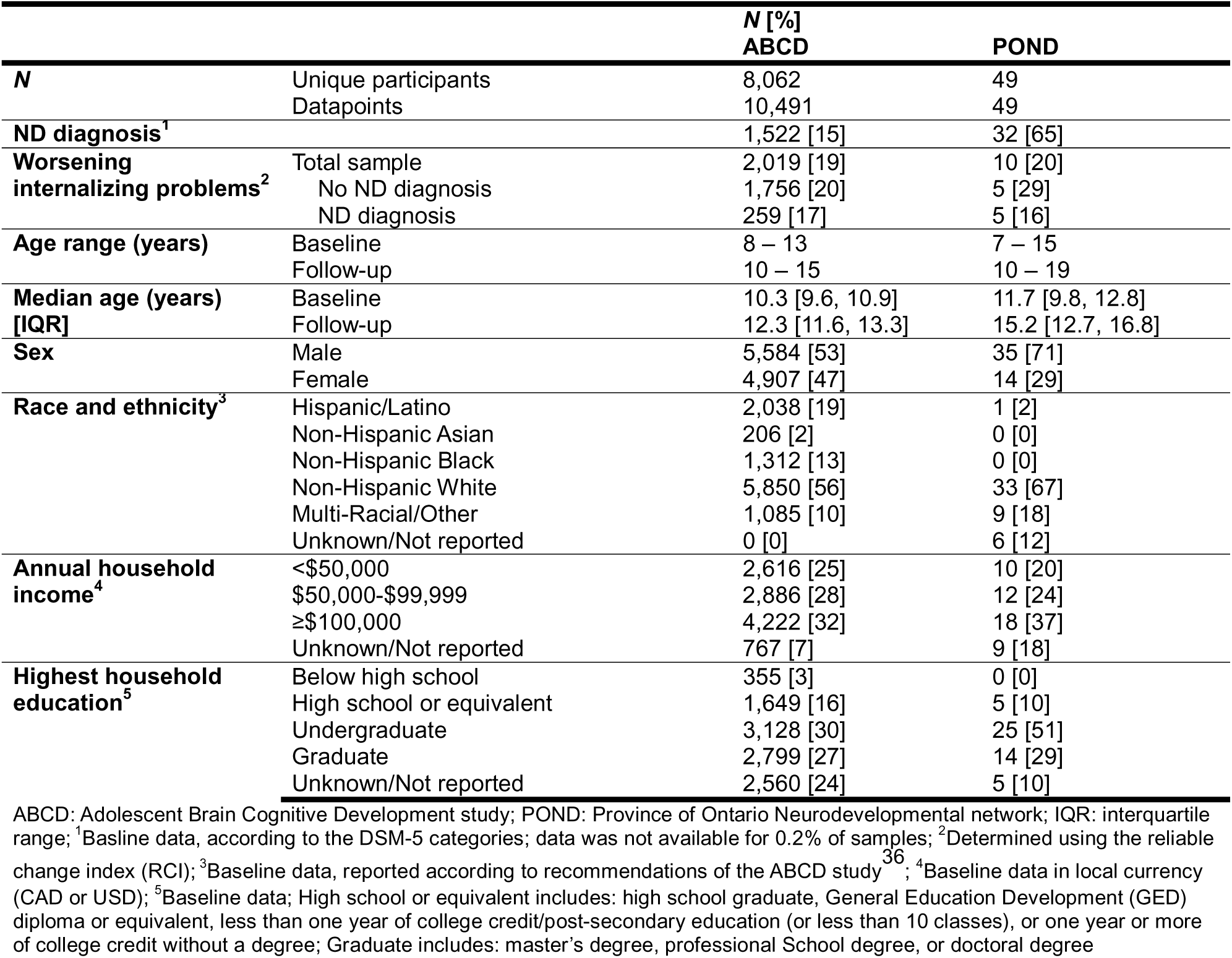
Participant demographics for the longitudinal analysis.

### Predicting cross-sectional internalizing problems

We first examined the features of brain structure that contribute to internalizing symptoms by predicting clinically significant (CBCL *T*-score ≥64) versus non-significant (CBCL *T*-score <64) problems cross-sectionally. The ROC curves across the entire sample and disaggregated by ND diagnosis are shown in **Figure 1A**. Across the cross-validation folds, the overall AUC was 0.80 (95% CI: [0.71, 0.88]), while the AUC for individuals with and without a diagnosis of an ND was 0.71 [0.60, 0.82] and 0.78 [0.68, 0.89], respectively. At the optimal prediction score cut-off (0.49 [0.46, 0.52]), an accuracy of 77% [68, 86] was achieved (ND: 62% [50, 75]; no ND: 82% [74, 90]). When only age and sex were included as predictors, AUC was reduced to 0.58 [0.56, 0.59].

**Figure 1:**
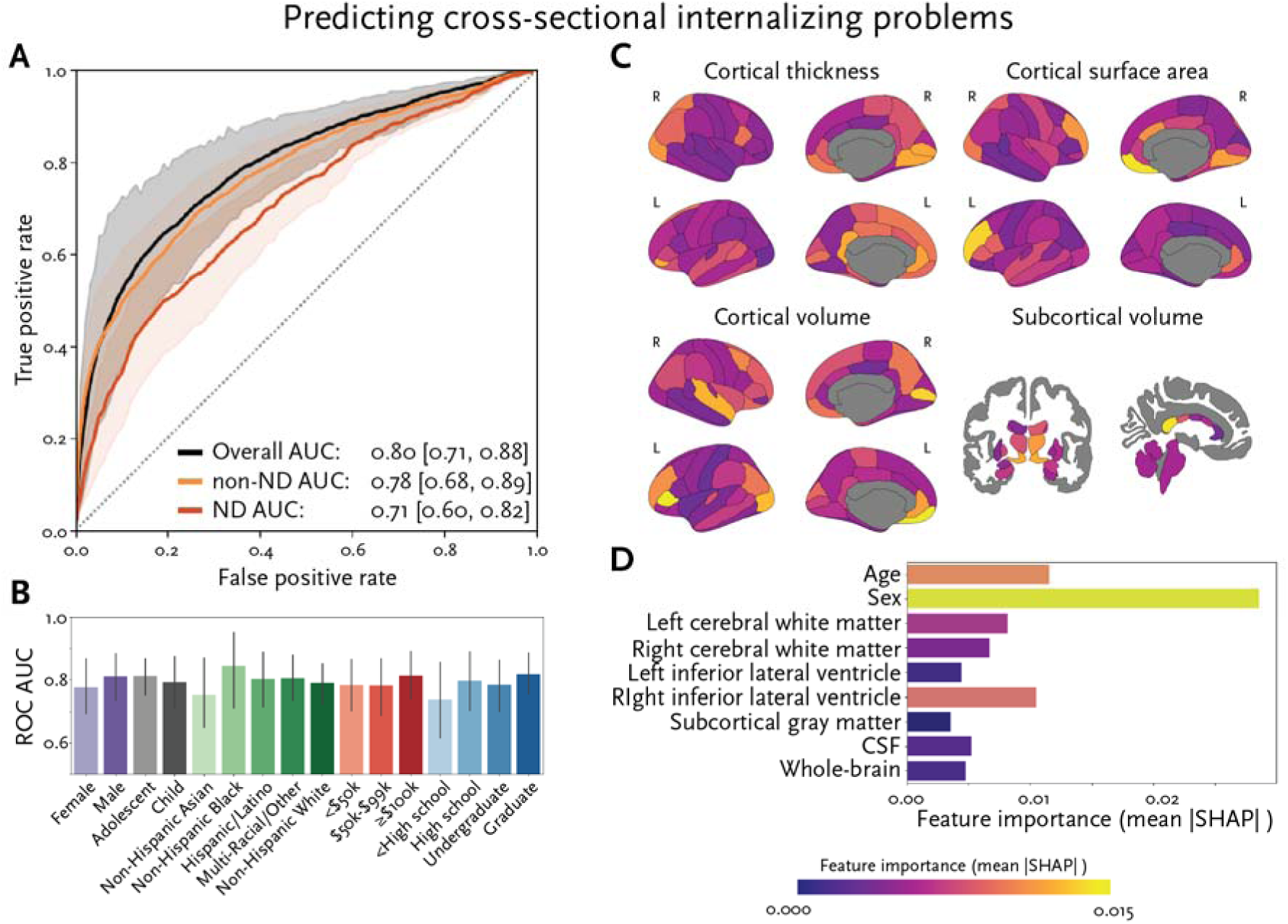
Performance and feature importances for the model predicting cross-sectional internalizing problems. (A) The mean [95% CI] receiving operating characteristic curve and corresponding area under the curve (AUC) across all data (black) and stratified by individuals with (red) and without (orange) an ND diagnosis. (B) AUCs in different sociodemographic groups (age, sex, race/ethnicity, annual household income, and highest household education level). (C) Feature importances (mean absolute SHAP values) for the cortical and subcortical features used in the prediction model. (D) Feature importances (mean absolute SHAP values) for the sociodemographic and global brain features.

AUCs were also compared amongst sociodemographic groups (age, sex, race/ethnicity, annual household income, and highest household education) to evaluate fairness (**Figure 1B**). No significant differences were observed (**Table 3**), indicating that our model performed comparably for the different sociodemographic groups.

**Table 3:**
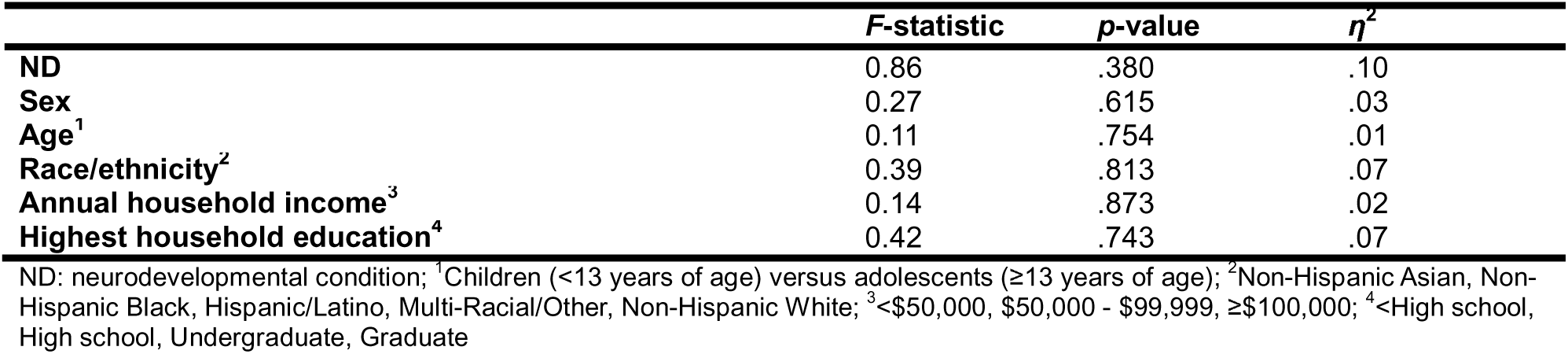
Statistics comparing AUC amongst sociodemographic groups for the cross-sectional prediction model.

| | *F*-statistic | *p*-value | $\eta^2$ |
| --- | --- | --- | --- |
| ND | 0.86 | .380 | .10 |
| Sex | 0.27 | .615 | .03 |
| Age<sup>1</sup> | 0.11 | .754 | .01 |
| Race/ethnicity<sup>2</sup> | 0.39 | .813 | .07 |
| Annual household income<sup>3</sup> | 0.14 | .873 | .02 |
| Highest household education<sup>4</sup> | 0.42 | .743 | .07 |

ND: neurodevelopmental condition; <sup>1</sup>Children (<13 years of age) versus adolescents (≥13 years of age); <sup>2</sup>Non-Hispanic Asian, Non-Hispanic Black, Hispanic/Latino, Multi-Racial/Other, Non-Hispanic White; <sup>3</sup><\$50,000, \$50,000 - \$99,999, ≥\$100,000; <sup>4</sup><High school, High school, Undergraduate, Graduate

Feature importances, which capture the relative extent to which each feature influences the final prediction, were calculated using the magnitude of SHAP values (**Figure 1C** and **D**). The most important features were sex (with females having a negative impact on prediction scores), the area and thickness of the right frontal pole, the areas of the left temporal pole, left rostral middle frontal, and right medial orbitofrontal gyri, and the volumes of the left medial orbitofrontal gyrus, left pars triangularis, right pericalcarine cortex, and posterior corpus callosum.

### Predicting worsening trajectories of internalizing problems

We then predicted whether an individual would experience worsening (RCI>1.96) versus non-worsening (RCI≤1.96) internalizing problems from baseline measures of brain structure. The ROC curves across the entire sample and disaggregated by ND diagnosis are shown in **Figure 2A**. Across the cross-validation folds, the AUC was 0.66 (95% CI: [0.65, 0.67]) for ABCD, while the AUC for POND was 0.80 [0.78, 0.81]. Stratified by ND diagnosis across both datasets, the AUCs for individuals with and without an ND were 0.73 (95% CI: [0.70, 0.76]) and 0.65 (95% CI: [0.65, 0.67]), respectively. At the optimal prediction score cut-off (0.53 [0.50, 0.55]), an accuracy of 71% [65%, 77%] was achieved (ABCD: 63% [58, 69]; POND: 79% [76, 73]), and higher accuracy was achieved in those with (73% [70, 76]) an ND compared to those without (65% [64, 66]).

**Figure 2:**
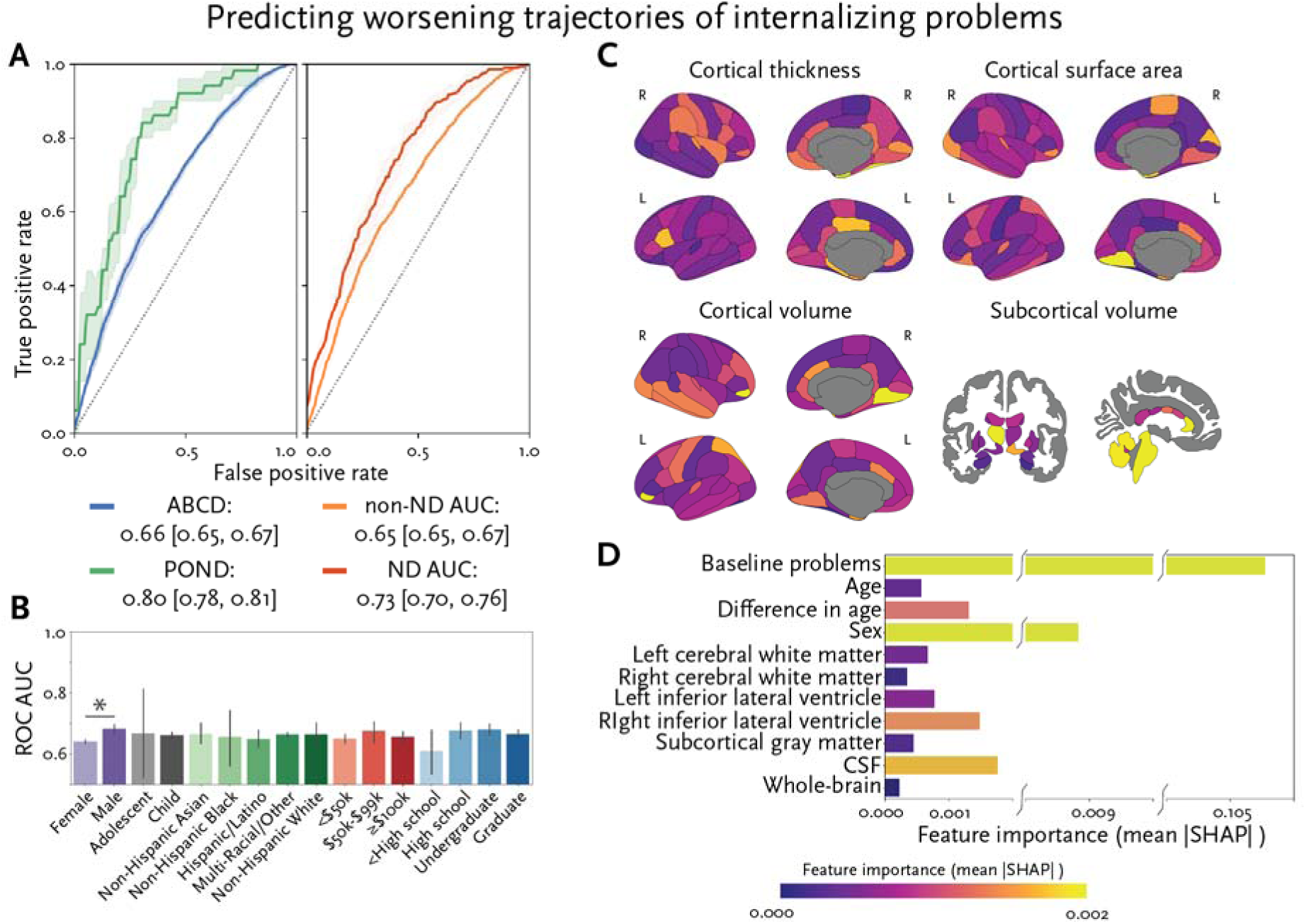
Performance and feature importances for the model predicting worsening trajectories of internalizing problems. (A) The mean [95% CI] receiving operating characteristic curve and corresponding area under the curve (AUC) across all data from ABCD (left; blue) and POND (left; green) and stratified by individuals with (right; red) and without (right; orange) an ND diagnosis. (B) AUCs in different sociodemographic groups (age, sex, race/ethnicity, annual household income, and highest household education level). (C) Feature importances (mean absolute SHAP values) for the cortical and subcortical features used in the prediction model. (D) Feature importances (mean absolute SHAP values) for the sociodemographic and global brain features.

AUCs were also compared amongst sociodemographic groups (age, sex, race/ethnicity, annual household income, and highest household education) to evaluate fairness (**Figure 2B**); a significant difference in sex was observed (*F*(1,8)=24.9, *p*=.001, *η*^2^=.76), with males having higher AUC compared to females (**Table 3**).

**Table 4:** Statistics comparing AUC amongst sociodemographic groups for the longitudinal prediction model.

| | *F*-statistic | *p*-value | $\eta^2$ |
| --- | --- | --- | --- |
| ND | 26.7 | .001 | .77 |
| Sex | 24.9 | .001 | .76 |
| Age<sup>1</sup> | 0.01 | .943 | .00 |
| Race/ethnicity<sup>2</sup> | 0.07 | .990 | .01 |
| Annual household income<sup>3</sup> | 1.03 | .387 | .15 |
| Highest household education<sup>4</sup> | 2.10 | .141 | .28 |

ND: neurodevelopmental condition; <sup>1</sup>Children (<13 years of age) versus adolescents (≥13 years of age); <sup>2</sup>Non-Hispanic Asian, Non-Hispanic Black, Hispanic/Latino, Multi-Racial/Other, Non-Hispanic White; <sup>3</sup><\$50,000, \$50,000 - \$99,999, ≥\$100,000; <sup>4</sup><High school, High school, Undergraduate, Graduate

Feature importances were calculated using the magnitude of SHAP values (**Figure 2C** and **D**). The most important features were the baseline score of internalizing problems (with fewer problems having a positive impact on prediction scores), sex (with females having a positive impact), the thicknesses of the right entorhinal cortex and fusiform, the areas of the right temporal pole and left lingual gyrus, and the volumes of the left thalamus, right lingual gyrus, anterior corpus callosum, and brainstem. Feature importance values from the longitudinal analysis were not significantly correlated with those from the cross-sectional analysis (*R*=0.02, *p*=0.816).

## Discussion

This study examined the utility of using deep learning to predict mental health problems from measures of brain structure. Cross-sectionally, good performance (mean AUC=0.80) was achieved across a large sample (*N*=14,523) of children and youth, providing insight into the neurobiology underlying internalizing problems beyond what could be revealed by traditional regression. Longitudinally (*N*=10,540), performance was sub-optimal for predicting worsening trajectories of internalizing problems in children and youth from the general population (mean AUC=0.66). However, the model performed well for an external testing set comprised primarily of youth with an ND (mean AUC=0.80), and when only considering individuals across the whole sample with an ND diagnosis (mean AUC=0.73). These results indicate that deep learning with measures of neurobiology could potentially improve our ability to detect which individuals, especially which neurodivergent individuals, are at risk of experiencing worsening trajectories of mental health, allowing for timely interventions to help mitigate the rising burden on youth, families, and healthcare systems around the world.

Deep learning was able to predict internalizing problems from cross-sectional measures of neurobiology with good performance (mean AUC=0.80). While cross-sectional prediction does not have direct clinical relevance, it does allow for the comparison of the utility of using symptom measures, rather than diagnostic labels, as prediction targets. As the popularity of artificial intelligence has grown, there has been substantial work on its application to predicting ND conditions, such as autism and attention-deficit/hyperactivity disorder, from measures of brain structure. However, despite having distinct behaviour-based diagnostic criteria, these ND conditions are highly heterogeneous in neurobiology and phenotype, and overlap with each other, and even typical development ^30,31^. This has posed two main challenges for the existing literature. First, for clinical translation, the observed heterogeneity necessitates larger sample sizes to capture the full spectrum of variability. Yet, most studies have used small sample sizes^37^. While the diagnostic prediction accuracies of these underpowered studies have been high when predicting ND diagnosis versus typical development, accuracy was shown to decrease with sample size, with the best powered studies only achieving an accuracy of just over 60%^37^. Second, considering the heterogeneity and overlap, there is evidence that the diagnostic labels do not align with the underlying neurobiology ^30,31^, which fundamentally limits work attempting to predict diagnosis from neurobiology. Here, we have instead predicted mental health symptoms experienced across neurotypical and neurodivergent children and youth from measures of brain structure. Across this sample, we have achieved an accuracy of 77%, a substantial improvement over the existing literature, supporting the use of transdiagnostic symptom measures as prediction targets.

Comparatively, the deep learning model trained to predict worsening trajectories of internalizing problems did not perform well for a sample of children and youth representative of the general population (ABCD; mean AUC=0.66). However, when applied to a small withheld external testing set comprised primarily of children with an ND diagnosis (POND; *N*=49), performance improved considerably (mean AUC=0.80). While this external sample was small, this provides some indication that the model holds value for predicting worsening symptoms in individuals at high-risk of mental health symptoms. When examining performance across the individuals with an ND diagnosis across the entire sample, performance improved considerably (mean AUC=0.73). This improvement cannot be explained by an increased representation of worsening changes in the ND group, as prevalence rates were comparable in individuals with (17%) and without (20%) an ND diagnosis. The performance difference was also not observed in the cross-sectional model, and the lack of significant difference in model performance across demographic variables (aside from sex, discussed later) suggests that sample characteristics alone are unlikely to explain this longitudinal-specific effect. Nonetheless, we acknowledge that unmeasured differences between the cross-sectional and longitudinal samples may have contributed to this pattern.

We hypothesize that biological differences associated with NDs more reliably predict subsequent mental health difficulties, potentially suggesting that mental health challenges may be part of the constellation of biological atypicality seen in NDs. There are ongoing debates about whether mental health difficulties in NDs represent shared or distinct mechanisms compared to the general population, with symptom overlap, measurement challenges, and diagnostic ambiguity obscuring the relationship between diagnosis and co-occurring mental health conditions^21,38^. While our findings point to the neurobiological signatures of mental health trajectories being more distinguishable in NDs, it remains unclear whether these reflect shared mechanisms or unique pathways intrinsic to neurodevelopmental conditions. These complexities underscore the importances of developing tailored assessment approaches and clarifying underlying mechanisms to improve prediction and support personalized care. In addition, there may be more heterogeneity in the biological correlates of mental health trajectories in NDs; while feature variability can make initial learning challenging for a deep learning model, it can eventually lead to more general and robust performance. Finally, the increased prevalence of clinically significant internalizing problems in the NDs may be a contributing factor – worsening trajectories that extend from elevated baseline traits may have more distinct, or well-defined, neurobiological correlates compared to trajectories that arise from low baseline symptoms.

The cross-sectional model is also able to provide insight into the importance of individual neurobiological features for predicting internalizing problems. Nearly all of the most important features resided in the prefrontal cortex and included the medial orbitofrontal cortex, along with the temporal poles and corpus callosum. The role of these brain structures in internalizing disorders has been consistently documented^7,39,40^, reflecting their roles in emotion processing, regulation, and awareness. The prefrontal cortex, in particular, has often been linked more strongly to anxiety over other domains of internalizing symptoms, such as depression ^41–43^, suggesting that our model may be particularly sensitive to this dimension, though this remains speculative. Furthermore, while the deep learning model relied on features already known to be associated with internalizing problems to make predictions, we have provided further insight into the multivariate complexities in these associations. For example, the importance of the lateral prefrontal cortex was specific to its surface area and volume, not thickness, and the patterns of importances throughout the brain were largely hemisphere dependent. Traditional approaches, like regression, typically search for linear brain-behaviour associations within each brain region and structural modality in isolation. The capability of deep learning models to learn from high-dimensional data allows these models to capture intricate patterns and relationships across many variables that might be challenging for traditional statistical methods. This helps advance our understanding of the neural correlates of mental health problems, which can be used to generate hypotheses for future research.

There was some overlap between features important for the cross-sectional and worsening models, for example the temporal pole and corpus callosum; however, the patterns of feature importances were strikingly different between the two models. This finding suggests that there are neuroanatomical patterns that influence internalizing problems over time, and these patterns are distinct from those that arise from the presence of clinically significant problems. In addition to brain features, baseline levels of internalizing problems were also important for predicting worsening trajectories of internalizing problems. Caution should be used in interpreting this finding through a clinical lens, given the limited range of the CBCL and the use of the reliable change index to identify worsening trajectories. Thus, this likely reflects the fact that individuals with very few baseline problems are more likely to exhibit worsening trajectories than individuals with problems already at the upper limit of the scale.

Our findings suggest that individual differences in brain structure may encode predictive information about both current and future internalizing symptoms. This indicates that structural features are not merely correlates of mental health status but may reflect underlying neurobiological vulnerabilities or compensatory mechanisms. For example, if characteristics of the prefrontal cortex predict current clinical status, this reinforces its established role in emotion processing, regulation, and awareness. Conversely, if other regions are more predictive of longitudinal symptom changes, this may point to roles in resilience, disorder progression, or neural plasticity. The absence of predictive value in certain regions also contributes to our understanding by delineating areas less involved in symptom emergence or stability. More broadly, these findings support the notion that the brain’s structural architecture captures meaningful interindividual variability in mental health risk and trajectory, and they offer insight into regions that may be most relevant for early detection and intervention.

Sex was also one of the most import predictive features in both the cross-sectional and worsening models. In the cross-sectional model, the feature importances indicated that being male positively impacted the predicted probability of having internalizing problems, while being female had a negative impact. While internalizing problems have classically been reported as increased in females compared to males^44^, more recent work has reported no sex differences in both neurotypical and neurodivergent populations^45^. Considering this work, we hypothesize that positive predictive impact of males in the cross-sectional model may reflect the male preponderance in neurodivergence^29^, which is in turn associated with increased internalizing problems^21^. On the other hand, in the longitudinal model, being female positively impacted the probability of predicted worsening internalizing problems, while being male had a negative impact. This is in line with evidence showing that both neurotypical and neurodivergent females are more likely to exhibit worsening longitudinal trajectories of internalizing problems across adolescence^46^. Together, the present study supports the importance of sex as an influencing factor on internalizing problems. Importantly, this study only considered biological sex, and not gender. There is some evidence of differences in internalizing problems between gender identities (e.g., Herrmann et al. (2024)^47^), although work in this domain has been limited; there is also evidence that sex differences in internalizing problems may be caused, in part, by gender socialization, rather than biological sex^48^. While considering biological sex as a feature alongside neurobiology in the deep learning models, rather than regressing out its effects, is a step towards addressing its contribution to mental health; future work should consider measures of gender alongside measures of biological sex.

The present study provides data demonstrating that AI approaches using biological markers can be useful both at the cross-sectional level to uncover complex relationships between biology and phenotype that are not typically amenable to traditional statistical methods, as well as for predicting emerging clinically important phenotypes longitudinally. As such it is not an exhaustive search for brain-behaviour relationships during development in mental health. We only considered measures of brain structure and limited sociodemographic measures (age and sex) as predictors of cross-sectional and longitudinal internalizing problems. Other neurobiological measures, such as brain function and structural connectivity, alongside genetic, environmental, and broader sociodemographic variables, likely also hold predictive value. We also only examined internalizing problems as a prediction target, which are just one aspect of mental health; examining the utility of predicting sub-scores of internalizing problems (e.g., anxious/depressed, withdrawn/depressed, and somatic complaints), externalizing problems, or both problem domains, is an important next step. Furthermore, we measured internalizing problems using a caregiver report, which have been associated with response biases. Other limitations include that, due to the use of the ABCD dataset, the bulk of the sample’s age is in the early adolescent years; this is particularly true for the longitudinal analysis, where two of the four datasets (HBN, HCP-D) cannot be used. Future work should include more younger children and older youth to ensure generalizability across development. We also did not consider the onset of the COVID-19 pandemic.

In conclusion, this study demonstrated that deep learning can be used to predict internalizing problems cross-sectionally, with performance exceeding that of comparable studies predicting ND diagnosis. We also showed that baseline measures of brain structure can predict worsening trajectories of internalizing problems, especially for children and youth with an ND. This suggests that neurobiology may offer a promising path towards identifying biomarkers of mental health trajectories in populations most vulnerable to adverse outcomes. Ultimately this type of prediction can support the delivery of proactive care aimed at preventing the escalation of mental health difficulties. While this study establishes the feasibility of this approach, future research is needed to evaluate this approach in prospective designs and real-world settings. Consideration must also be given to the availability and integration of neuroimaging within current clinical pathways. To ensure acceptability, equity, and clinical relevance, co-design with both clinicians and the neurodivergent community will be essential for translating these tools into practice. This work adds to the growing evidence that responsible applications of AI can advance personalized approaches to mental health care.

## Acknowledgements

Funding for the current study was provided by the Canadian Institutes of Health Research and New Frontiers in Research Fund.

## ABCD

Data used in the preparation of this article were obtained from the Adolescent Brain Cognitive Development^SM^ (ABCD) Study (https://abcdstudy.org), held in the NIMH Data Archive (NDA). This is a multisite, longitudinal study designed to recruit more than 10,000 children age 9-10 and follow them over 10 years into early adulthood. The ABCD Study® is supported by the National Institutes of Health and additional federal partners under award numbers U01DA041048, U01DA050989, U01DA051016, U01DA041022, U01DA051018, U01DA051037, U01DA050987, U01DA041174, U01DA041106, U01DA041117, U01DA041028, U01DA041134, U01DA050988, U01DA051039, U01DA041156, U01DA041025, U01DA041120, U01DA051038, U01DA041148, U01DA041093, U01DA041089, U24DA041123, U24DA041147. A full list of supporters is available at https://abcdstudy.org/federal-partners.html. A listing of participating sites and a complete listing of the study investigators can be found at https://abcdstudy.org/consortium_members/. ABCD consortium investigators designed and implemented the study and/or provided data but did not necessarily participate in the analysis or writing of this report. This manuscript reflects the views of the authors and may not reflect the opinions or views of the NIH or ABCD consortium investigators. The ABCD data repository grows and changes over time. The ABCD data used in this report came from DOI: 10.15154/z563-zd24. DOIs can be found at https://nda.nih.gov/abcd.

## HBN

This manuscript was prepared using a limited access dataset obtained from the Child Mind Institute Biobank, the Healthy Brain Network (HBN). This manuscript reflects the views of the authors and does not necessarily reflect the opinions or views of the Child Mind Institute.

## HCP-D

Research reported in this publication was supported by the National Institute of Mental Health of the National Institutes of Health under Award Number U01MH109589 and by funds provided by the McDonnell Center for Systems Neuroscience at Washington University in St. Louis. The HCP-Development 2.0 Release data used in this report came from DOI: 10.15154/1520708.

## POND

This research was conducted with the support of the Ontario Brain Institute (POND, PIs: Anagnostou/Lerch), an independent non-profit corporation, funded partially by the Ontario government. The opinions, results and conclusions are those of the authors and no endorsement by the Ontario Brain Institute is intended or should be inferred.

## Conflicts of interest

AK has a patent for holly^TM^ (formerly Anxiety Meter) with royalties paid from Awake Labs. AK has received consulting fees from DNAStack and Shaftesbury. EA has received grants from Roche and Anavex, served as a consultant to Roche, Quadrant Therapeutics, Ono, and Impel Pharmaceuticals, has received in-kind support from AMO Pharma and CRA-Simons Foundation, received royalties from APPI and Springer, received an editorial honorarium from Wiley, and has a patent for holly^TM^ (formerly Anxiety Meter). The remaining authors have no potential conflicts of interest to report.

## Data and code availability

Access to the data used in this study is controlled by each of the datasets (ABCD, HBN, HCP-D, and POND). Details on the data usage agreements can be found in the **Supplementary material**. Code is available on GitHub (https://github.com/marlvan/internalizing_prediction).

## Author contributions

Conceptualization: MMV, AK; Data curation: MMV, BS; Formal analysis: MMV; Funding acquisition: MMV, JPL, EA, AK; Investigation: MMV; Methodology: MMV; Project administration: AK; Resources: EK, JJ, MA, PDA, JC, RJS, MJT, JPL, EA, AK; Software: MMV; Supervision: AK; Validation: MMV; Visualization: MMV; Writing – original draft: MMV; Writing – review & editing: MMV, BS, NB, AA, EK, JJ, MA, AI, PDA, JC, RJS, MJT, JPL, EA, AK

## Supplementary material

### Recruitment strategies

The ABCD study is a population-based cohort study that aimed to recruit and follow 11,500 children that were representative of the population of the United States; full details on recruitment are described elsewhere^49^. Briefly, children between 9-10 years of age were recruited through public and private elementary schools to 21 nationally distributed sites, whose catchment areas contained one in every five eligible children in the US. To minimize bias and ensure a representative sample, school selection was informed by sociodemographic factors and recruitment was dynamically monitored. After their initial recruitment, participants were asked to return every two years for imaging and other assessments. Children were excluded for a lack of English fluency, neuroimaging contraindications, a major neurological disorder, <28 weeks gestational age, <1,200g birth weight, >1 month hospitalization due to birth complications, uncorrected sensory impairments, a current diagnosis of schizophrenia, autism, intellectual disability, or alcohol/substance use disorder, a history of traumatic brain injury, or an unwillingness to complete the testing protocol^50,51^.

The objective of HBN was to create a large transdiagnostic dataset that captures a broad range of clinical developmental psychopathology^23^. A community-referred recruitment strategy was used, where advertisements that targeted families with concerns about their child’s psychiatric well-being were distributed to community members, educators, and care providers in the New York City area of the United States. Children and youth were eligible for HBN if they were between 5 – 21 years of age, were fluent in English, could provide verbal assent, and whose parent/caregiver could provide informed consent. Exclusion criteria included the presence of a debilitating neurological disorder that prevented participation in the full protocol, injury or disease-related acute encephalopathy, a known neurodegenerative disorder, uncorrected sensory impairments, a diagnosis of schizophrenia, schizoaffective disorder, or bipolar disorder given in the past six months, currently untreated suicidality or homicidality with onset within the past three months, a history of substance dependence that requires chemical replacement therapy, and intoxication at any study visit.

The HCP-D study is a sub-study of the Human Connectome Project, which aims to characterize the development of brain connectivity^24^. Children and adolescents between 5 – 21 years of age were recruited from four cities in the United States (Boston, Los Angeles, Minneapolis, and St. Louis). Participants were excluded based on insufficient English fluency, premature birth, a lifetime history of serious medical or endocrine conditions, treatment for a serious medical problem, treatment for more than one year by a specialist for certain disorders, head injury, hospitalization for at least two days for certain conditions, receiving special education services at school, and contraindications for neuroimaging.

The objective of the POND study is to better understand the biological underpinnings of neurodevelopmental conditions and consists of children and adolescents with and without these conditions. Participants are recruited from clinical research sites in four cities on Ontario, Canada, with three institutions collecting neuroimaging data. POND contains a longitudinal component, where participants were asked to return at least two years after their first visit. Participants were recruited if they had a clinical diagnosis of a neurodevelopmental condition, including autism, attention-deficit/hyperactivity disorder, and obsessive-compulsive disorder or typically developing controls. Typically developing participants were recruited using flyers posted in the community and word-of-mouth and were excluded on the basis of a diagnosed neurodevelopmental, psychiatry, or neurological condition, a first-degree family member with a diagnosed neurodevelopmental condition, or premature birth. All participants were screened for contraindications for neuroimaging and sufficient English for consent and the testing protocols.

### Assent/consent procedures

For the ABCD study, parent(s)/caregiver(s) provided written informed consent for their child’s participation, while written assent was obtained from the participant^52^. For HBN, participants 18 or older provided written informed consent, while those under 18 provided verbal assent and written informed consent was obtained by a parent/caregiver^23^. For the HCP-D study, parent(s)/caregiver(s) provided written informed consent for their child’s participation. For the POND study, written informed consent was obtained from participants who had the capacity to do so; otherwise, written consent was obtained from a parent(s)/caregiver(s), and verbal assent was obtained from the participant, as per institutional ethics board SOPs.

### Institutional review board approvals

The majority of ABCD’s 21 sites used a centralized Institutional Review Board at the University of California, San Diego which reviewed and approved the study’s protocol, with some sites having used a local Institutional Review Board^53^. The study protocol for HBN was reviewed and approved by the Chesapeake Institutional Review Board. For the HCP-D study, a centralized Institutional Review Board at Washington University, St. Louis reviewed and approved the study’s protocol. For the POND study, each site’s Institutional Review Board approved the study protocol.

### Data usage agreements

Access to the ABCD and HCP-D data was obtained under the NIMH Data Archive Data Use Agreement #15619 and #18618 for the Adolescent Brain Cognitive Development (ABCD)/Connectome Coordination Facility (CCF) permission group.

Access to the HBN phenotypic data was granted after approval of a data usage agreement awarded to A. Kushki; an agreement was not required to access the imaging data. For the POND data, access is available as the study team reflected in the authorship line is also part of the POND study team, according to internal procedures.

### Sociodemographic assessments

Biological sex was collected as part of all four study protocols by asking parent(s)/caregiver(s) to specify what sex the child was assigned at birth. For ABCD, options included male, female, intersex male, and intersex female. For HCP-D, male, female and other were options. For POND and HBN, only male and female were options.

Race and ethnicity data was collected as part of all study protocols. The ABCD study provides the data pre-categorized into five recommended distinct racial/ethnic categories (Hispanic/Latino, non-Hispanic Asian, non-Hispanic Black, non-Hispanic White, and Multi-Racial/Other)^36^, after asking the parent(s)/caregiver(s) to specify what race they consider their child to be. The recommended procedure was followed to categorize the raw race/ethnicity data from the remaining studies (described below) into these five categories.

In HBN, parent(s)/caregiver(s) are asked to select one category for their child’s ethnicity (not Hispanic or Latino, Hispanic or Latino, decline to specify, or unknown) and race (White/Caucasian, Black/African American, Hispanic, Asian, Indian, Native American Indian, American Indian/Alaskan Native, Native Hawaiian/Other Pacific Islander, two or more races, or other race). In HCP-D, parent(s)/caregiver(s) or participants were asked to specify their ethnicity (Hispanic or Latino, Not Hispanic or Latino, or unknown/not reported), and asked what race they considered their child/themselves to be (American Indian/Alaska Native, Asian, Hawaiian or Pacific Islander, Black or African American, White, more than one race, and/or unknown or not reported); multiple categories were mapped to the “more than one race” category. In POND, parent(s)/caregiver(s) were asked to select all racial/ethnic groups that applied to their child (Aboriginal, Arab, Black, Chinese, East Asian, Filipino, Japanese, Jewish, Korean, Latin American/Hispanic, South Asian, Southeast Asian, West Asian, White, other, don’t know, prefer not to answer).

Annual household income and highest household education were used as measures of socioeconomic status. For ABCD, HBN, and POND, parent(s)/caregiver(s) were asked to select their annual household income from several provided income bands. For HCP-D, either the participant or their parent(s)/caregiver(s) provided the numerical value for their annual household income. This raw data was used to categorize participants into low (<$50,000), medium ($50,000-$99,999) and high (≥$100,000) income households, along with unknown/not reported. Note that although POND is a Canadian study, income levels were not adjusted for the exchange rate. The highest education level achieved by one or two parent(s)/caregiver(s), where appropriate, was collected in all four datasets by asking responders to select from a provided list. This raw data was used to categorize the education levels for each parent/caregiver into several ordinal categories: below high school, high school or equivalent, Associate/Bachelor’s degree, and unknown/not reported. High school or equivalent includes high school graduate, general education development (GED) diploma or equivalent, and any college education that did not result in a degree. Graduate degree includes degrees from master’s, doctoral, and professional school educations. When data from two caregivers was available, the highest level of education was selected.

ND diagnoses were also available for ABCD, HBN, and POND; for HCP-D, participants were excluded from the study based on the presence of a clinical diagnosis. For the purposes of this study, we defined an ND according to the diagnoses in the fifth edition of the Diagnostic and Statistical Manual of Mental Disorders (DSM-5)^54^: intellectual disabilities, communication disorders, autism spectrum disorder, attention-deficit/hyperactivity disorder, specific learning disorder, motor disorders, and other neurodevelopmental disorders. For ABCD, parents and youth completed a computerized version of the Kiddie Schedule for Affective Disorders and Schizophrenia (K-SADS) for DSM-5 (K-SADS-COMP). Across both parent and youth reports, if the K-SADS indicated a diagnosis in any of the neurodevelopmental conditions (past, present, or in partial remission), the participant was labeled as having a neurodevelopmental condition. In HBN, after all assessments are complete, DSM-5 diagnoses that reach a consensus across clinicians are assigned; these diagnoses were used to categorize the participants as having a neurodevelopmental condition. In POND, diagnostic assessments and expert clinical judgement (e.g., ^55–58)^ are used to assign a label of typical development or assign a primary diagnosis of a neurodevelopmental and related conditions (e.g., obsessive-compulsive disorder), which was used to determine the presence of a neurodevelopmental condition. In the cross-sectional sample, 84% of participants with any ND diagnosis had attention-deficit/hyperactivity disorder, 14% had autism, 7% had communication disorders, 1% had intellectual disabilities, 4% had motor disorders, 12% had specific learning disorders, and <1% had other neurodevelopmental disorders.

### Reliable change index

The reliable change index (RCI ^26^) was used to quantify whether a change in internalizing problems between two timepoints was larger than what could have been solely attributed to measurement error. We selected the Reliable Change Index (RCI) because it accounts for measurement reliability, distinguishes true clinical change from error, and is widely validated in clinical research. Alternatives such as percent change, the Standard Deviation Index, discrepancy scores, RCI with practice effects, and minimum important difference either ignore reliability or require additional data not available in our study. The index is computed by dividing the difference between the participant’s problem scores by the standard error associated with the assessment. Mathematically:

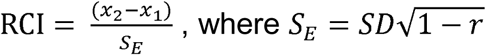

Here, *x*_1_ and *x*_2_ are the scores for the two timepoints, and *SE*_E_ is the standard error of the score, which is computed using the scores standard deviation (*SD*=10) and reliability (*r*=0.90^25^). This formula produces a *z*-score, and thus RCIs exceeding ±1.96 indicate a statistically significant, or reliable, change at the 5% threshold for significance. This cutoff was used to categorize participants as having worsening (RCI>1.96), improving (RCI<-1.96), or stable (−1.96≤RCI≤1.96) trajectories of internalizing problems.

### Imaging protocols

T1-weighted images were obtained as part of the imaging protocols for all four studies. For ABCD, data was collected from the 21 sites using three types of 3T scanners, whose acquisition parameters are presented in **Supplemental Table 1**. HBN collected data from four sites using one 1.5T and three 3T scanners, two of which used the same acquisition parameter (**Supplemental Table 2**). All four sites that collected data for HCP-D used the same Siemens 3T Prisma scanner, and the acquisition parameters are presented in **Supplemental Table 3**. For the POND study, data was collected using two 3T Siemens scanners at three imaging sites (**Supplemental Table 4**).

**Supplemental Table 1:** T1-weighted acquisition parameters for the ABCD study.

|  | Siemens<br>Prisma | Philips<br>Achieve dStream<br>Ingenia | GE<br>Discovery |
| --- | --- | --- | --- |
| **Matrix** | 256x256 | 256x256 | 256x256 |
| **Slices** | 176 | 225 | 208 |
| **FOV** | 256x256 | 256x256 | 256x256 |
| % FOV phase | 100 | 93.75 | 100 |
| Resolution (mm) | 1.0×1.0×1.0 | 1.0×1.0×1.0 | 1.0×1.0×1.0 |
| TR (ms) | 2500 | 6.31 | 2500 |
| TE (ms) | 2.88 | 2.90 | 2.00 |
| TI (ms) | 1060 | 1060 | 1060 |
| Flip angle (°) | 8 | 8 | 8 |
| Parallel imaging | 2x | 1.5×2.2 | 2x |
| Acquisition time (min:sec) | 7:12 | 5:38 | 6:09 |

**Supplemental Table 2:** T1-weighted acquisition parameters for the HBN study.

|  | Siemens Avanto | Siemens Trio/Prisma |
| --- | --- | --- |
| Matrix | 256×256 | 320×320 |
| Slices | 176 | 224 |
| FOV | 256×256 | 256×256 |
| % FOV phase | 100 | 100 |
| Resolution (mm) | 1.0×1.0×1.0 | 0.8×0.8×0.8 |
| TR (ms) | 2730 | 2500 |
| TE (ms) | 1.64 | 3.15 |
| TI (ms) | 1000 | 1060 |
| Flip angle (°) | 7 | 8 |
| Parallel imaging | 2x | 2x |
| Acquisition time (min:sec) | 6:32 | 7:19 |

**Supplemental Table 3:** T1-weighted acquisition parameters for the HCP-D study.

|  | Siemens Prisma |
| --- | --- |
| Matrix | 320×320 |
| Slices | 208 |
| FOV | 256×256 |
| % FOV phase | 93.8 |
| Resolution (mm) | 0.8×0.8×0.8 |
| TR (ms) | 2500 |
| TE (ms) | 1.81/3.6/5.39/7.18 |
| TI (ms) | 1000 |
| Flip angle (°) | 8 |
| Parallel imaging | 2 |
| Acquisition time (min:sec) | 8:22 |

**Supplemental Table 4:**
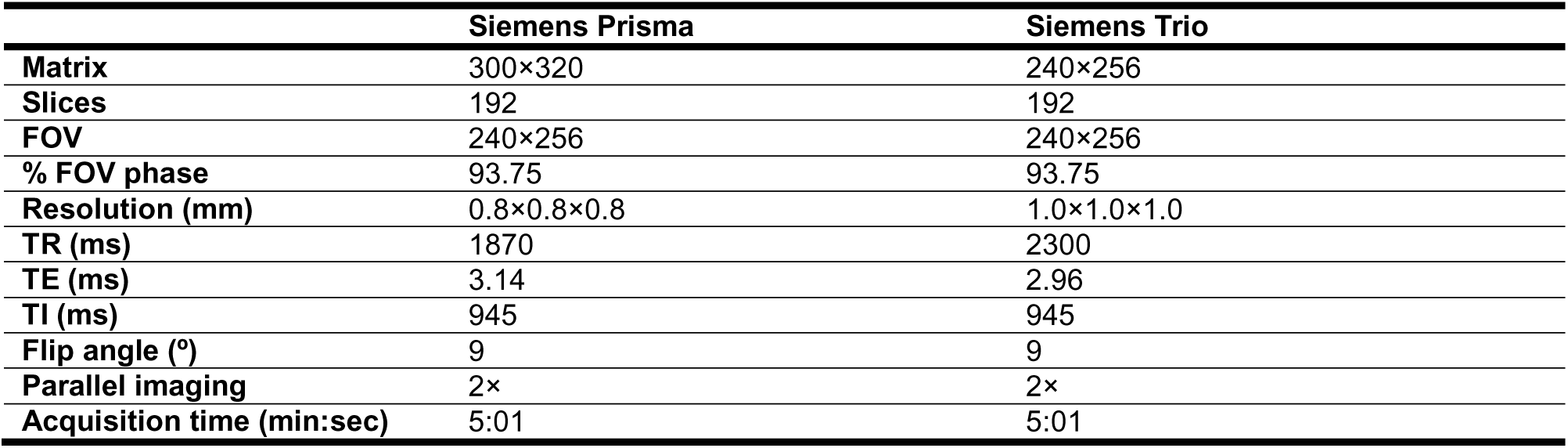
T1-weighted acquisition parameters for the POND study.

### Imaging preprocessing

For all datasets, the FreeSurfer image analysis suite^27^ was used to perform cortical reconstruction and volume segmentation of the structural MRIs. Measures of cortical thickness, cortical surface area, and cortical volume of the 68 region Desikan-Killiany parcellation^16^ were extracted as features from the FreeSurfer outputs, alongside the volumes provided by the subcortical segmentation^59^, which includes the subcortical structures, ventricles, white matter, brainstem, cerebellum, and whole-brain. For the ABCD study, the data was tabulated by the ABCD Data Analysis and Informatics Center using FreeSurfer version 5.3 or version 6.0, which was modified to account for intensity scaling and inhomogeneity correction that were already applied to the T1-weighted images^60^. For HCP-D, the Lifespan HCP Release 2.0 includes data processed using the HCP Pipelines (version 4.0.0^61^) which includes a modified version of FreeSurfer (version 6.0), from which tabulated data was extracted. Data from POND and HBN was processed in-house through a FreeSurfer (version 5.3) pipeline. We include the FreeSurfer boilerplate text below for completeness.

Cortical reconstruction and volumetric segmentation was performed with the Freesurfer image analysis suite, which is documented and freely available for download online (http://surfer.nmr.mgh.harvard.edu/). The technical details of these procedures are described in prior publications^59,62–74^. Briefly, this processing includes motion correction and averaging^73^ of multiple volumetric T1 weighted images (when more than one is available), removal of non-brain tissue using a hybrid watershed/surface deformation procedure ^72^, automated Talairach transformation, segmentation of the subcortical white matter and deep gray matter volumetric structures^59,66^ (including hippocampus, amygdala, caudate, putamen, ventricles) intensity normalization^75^, tessellation of the gray matter white matter boundary, automated topology correction^65,76^, and surface deformation following intensity gradients to optimally place the gray/white and gray/cerebrospinal fluid borders at the location where the greatest shift in intensity defines the transition to the other tissue class^62–64^. Once the cortical models are complete, a number of deformable procedures can be performed for further data processing and analysis including surface inflation^67^, registration to a spherical atlas which is based on individual cortical folding patterns to match cortical geometry across subjects^68^, parcellation of the cerebral cortex into units with respect to gyral and sulcal structure^28,69^, and creation of a variety of surface based data including maps of curvature and sulcal depth. This method uses both intensity and continuity information from the entire three dimensional MR volume in segmentation and deformation procedures to produce representations of cortical thickness, calculated as the closest distance from the gray/white boundary to the gray/CSF boundary at each vertex on the tessellated surface^64^. The maps are created using spatial intensity gradients across tissue classes and are therefore not simply reliant on absolute signal intensity. The maps produced are not restricted to the voxel resolution of the original data thus are capable of detecting submillimeter differences between groups. Procedures for the measurement of cortical thickness have been validated against histological analysis ^77^ and manual measurements^78,79^. FreeSurfer morphometric procedures have been demonstrated to show good test-retest reliability across scanner manufacturers and across field strengths^70,74^.

The distribution of cortical morphological features can vary between the site/scanner from which it is collected^80^. However, we also observed effects of site/scanner on the CBCL internalizing scores. For example, for the cross-sectional analysis, a significant effect of scanner was observed for ABCD (*H*(31)=219.3, *p*<.001, *η*^2^=0.02) and POND (*H*(4)=36.4, *p*<.001, *η*^2^=0.06), likely reflecting the geographic diversity and site-specific recruitment strategies, respectively. While techniques such as ComBat harmonization can be used to adjust for site effects while preserving covariate effects such as the RCIs^80^, the algorithm can perform poorly when there is high collinearity between site and these covariates^81^. Thus, we chose not to correct for multisite effects.

### Quality control

The ABCD study provided measures relating to data quality^60^. Manual quality control was performed by trained technicians prior to the FreeSurfer pipeline to visually identify images with unacceptable severe artifacts or irregularities. After FreeSurfer, technicians also reviewed the quality of the extracted surfaces, and provided an overall quality control score which indicated whether the surfaces were recommended for inclusion or exclusion. Only participants who passed both aspects of quality control were retained for analyses. For HBN, T1-weighted images were manually inspected in-house for motion-related artifacts using a three-point scale, and those with severe motion artifacts were excluded. For HCP-D, only images rated highly (“good” or “excellent”) on a four-point scale by technicians are processed through the HCP preprocessing pipeline; thus, all FreeSurfer data available passed quality control. Data from the POND study was also processed through the CIVET image-processing pipeline^82^, which provides an automated index of quality control that was used to exclude participants for the analyses.

### Datapoint selection

A flowchart of the study sample is presented in **Supplemental Figure 1**. The number of datapoints is the total number of unique pairs of CBCL assessments and MR scans across all participants. While only one CBCL assessment was acquired per session in all four studies, some POND and HBN participants had multiple structural images acquired during a single scanning session. For both the cross-sectional and longitudinal analyses, if multiple structural images were eligible for the final sample, one was selected at random.

**Supplemental Figure 1:**
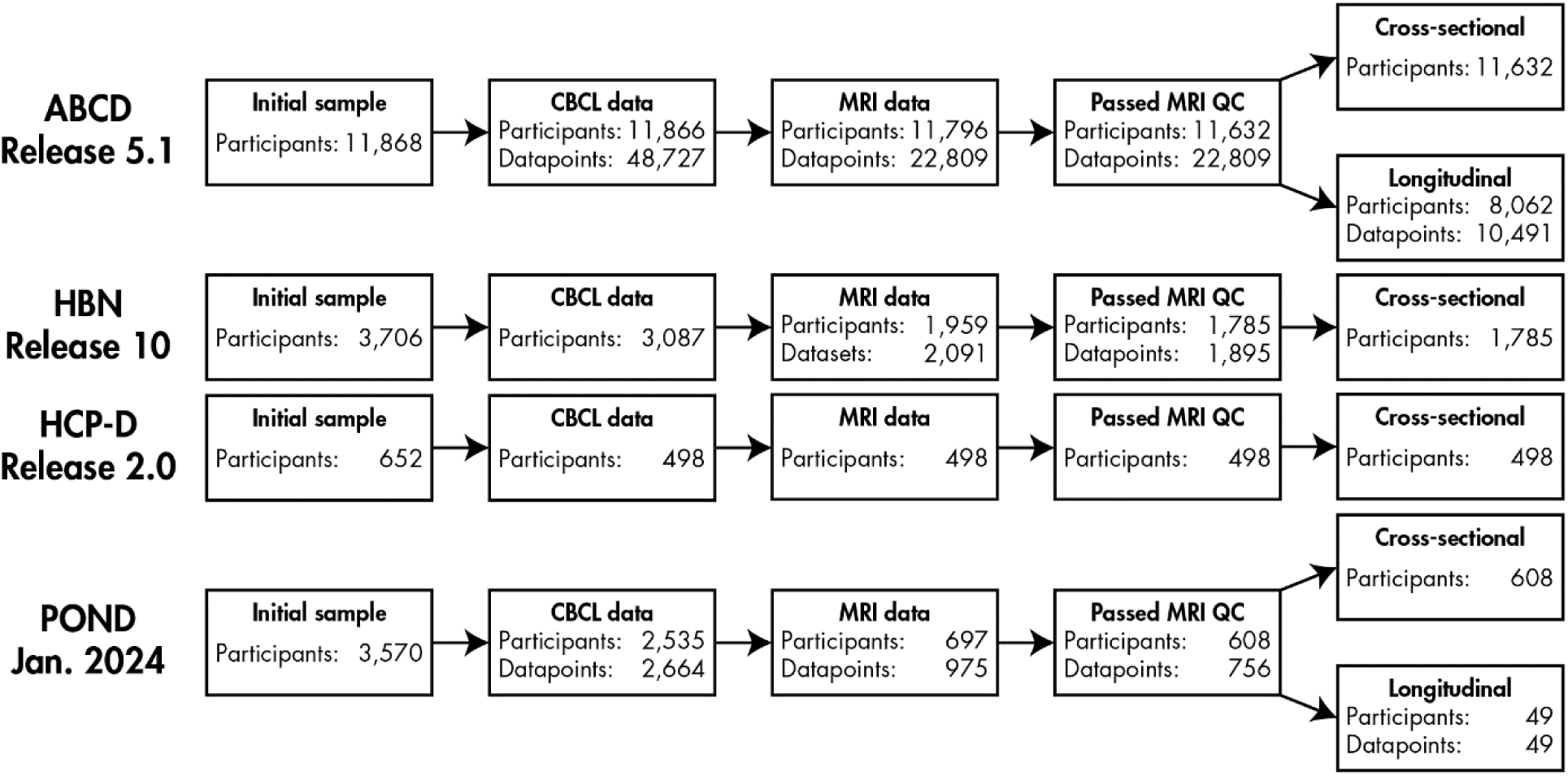
Flowchart of the study samples. For the cross-sectional analysis, one datapoint (CBCL assessment and structural image) was selected for each participant. The ABCD and POND datasets provided longitudinal data, from which pairs of datapoints were selected for the longitudinal analysis. ABCD: Adolescent Brain Cognitive Development study; HBN: Healthy Brain Network; HCP-D: Human Connectome Project Development study; POND: Province of Ontario Neurodevelopmental network; CBCL: Child Behaviour Checklist; MRI: magnetic resonance imaging; QC: quality control

For the cross-sectional analysis, a single session was selected from the longitudinal data available after quality control from ABCD and POND. For ABCD, given the higher proportion of participants with data from the initial recruitment session (9 – 11 years of age), the datapoint that gave the most uniform distribution across age was selected (per a Kolmogorov-Smirnov test). For POND, to minimize between-dataset differences in age, the session whose age best matched the distribution of the ABCD sample was selected.

For the longitudinal analysis, partial data was available for a third timepoint in the ABCD study. Thus, to maximize the total number of included datapoints, measures of brain structure from the initial assessment were used to predict internalizing problems at the first follow-up, and measures of brain structure from the first follow-up were used to predict internalizing problems at the second follow-up. For POND, if multiple longitudinal timepoints were available, the timepoints that best overlapped with the age range of the ABCD sample were selected. The distribution of the baseline and follow-up age is presented in **Supplemental Figure 2**.

**Supplemental Figure 2:**
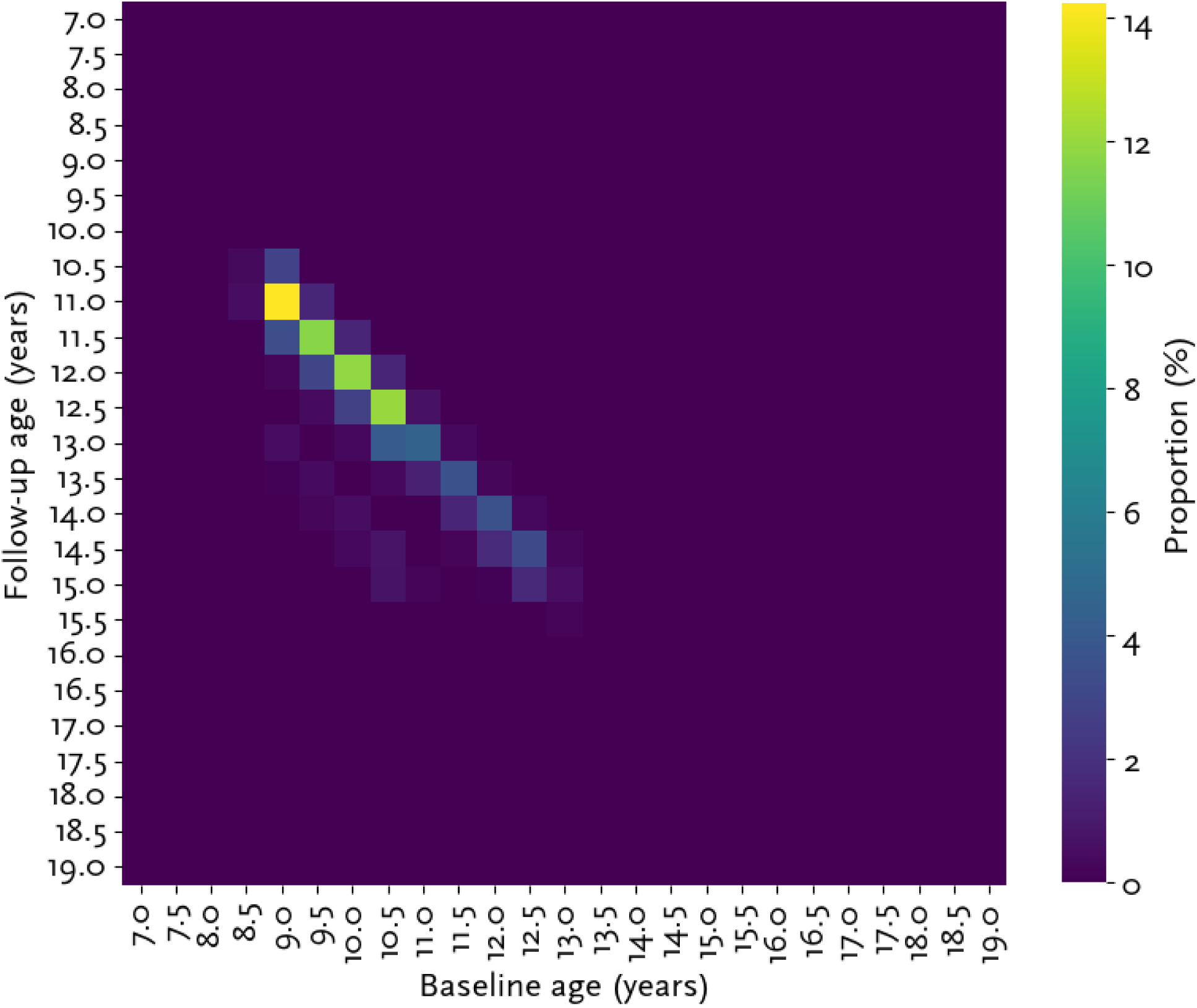
Age distribution at baseline and follow-up. For the longitudinal data, the proportion of participants (%) with each baseline and follow-up age combination is plotted, with warmer colours indicating a higher proportion.

### Deep learning model architecture

The model was implemented in Python (version 3.11) using the Keras framework (version 2.13.1;^83^) with TensorFlow (version 2.13.1;^84^) as the backend engine.

The model architecture was a multilayer perceptron, consisting of an input layer, blocks of hidden layers, and an output layer. Each block of hidden layers in the deep learning model consisted of a dense layer with rectified linear unit (RELU) activation, mitigating overfitting by applying L1 and L2 regularization to the layer’s kernel followed by a dropout layer. Sigmoid activation was used in the output layer to predict the probability of the positive class. The models were trained to minimize binary cross-entropy loss, using stochastic gradient descent with an adaptive moment estimation (ADAM) optimizer. Due to imbalances between the classes, the models were trained using a balanced batch generator, which over-samples the minority class to ensure balanced data within each training batch^85^.

A nested 5-fold stratified cross-validation scheme was used to tune, train, and test the model, with an inner loop for hyperparameter tuning and an outer loop for training and testing (**Supplemental Figure 3**). For the cross-sectional analysis, this scheme was performed using data aggregated across all four datasets. For the longitudinal analysis, given the imbalance between the POND and HBN datasets, the scheme was performed using ABCD, withholding POND as an external testing set.

**Supplemental Figure 3:**
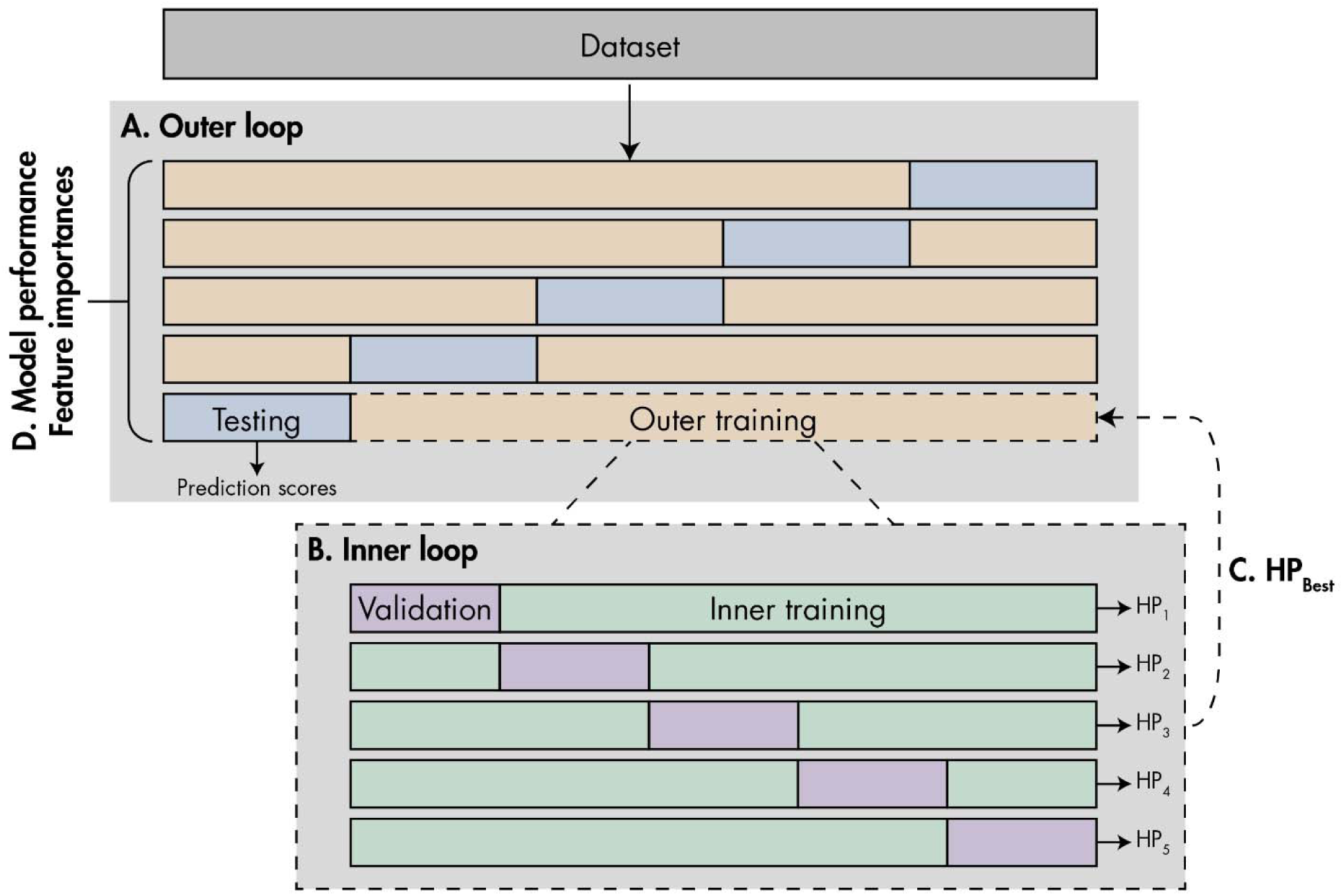
The nested cross-validation scheme used to train and test the deep learning models. The outer loop (A) uses 5-fold stratified cross-validation to split the data into testing (blue) and training (orange) sets. The inner loop (B) consists of a nested 5-fold stratified cross validation, where the outer training set is further split into validation (purple) and inner training (green) sets. Each validation fold is used to tune the model’s hyperparameters (HP), and the best performing hyperparameter set (C; HP_Best_) is selected to re-train the model using the entire outer training set, and prediction scores are calculated for the testing set. Measures of model performance and feature importances are then averaged across the outer folds (D).

The hyperparameter tuning was used to optimize the learning rate, batch size, and number of hidden blocks, as well as the number of neurons, regularization penalties, and dropout rate for each hidden block. Search criteria are presented in **Supplemental Table 5**. Given the class imbalance, the tuning was performed with the objective of maximizing the area under the precision-recall curve (AUC) of the validation set, using early stopping with a patience of 10 epochs and a maximum of 100 to prevent overfitting.

**Supplemental Table 5:** Search criteria for hyperparameter tuning.

|  |  | Values |  |  |
| --- | --- | --- | --- | --- |
| Learning rate |  | [0.1, 0.01, 0.001, 0.0001] |  |  |
| Batch size |  | [8, 16, 32, 64, 128, 256, 512, 1024] |  |  |
|  |  | Minimum | Maximum | Step size |
| Learning rate | | $1e^{-10}$ | 0.1 | Log sampling |
| Number of hidden blocks |  | 1 | 3 | 1 |
| Per hidden block | Number of neurons | 1 | $N_{\text{features}}$ | 1 |
|  | Dropout rate | 0.1 | 0.9 | 0.1 |
| | L1 regularization penalty | $1e^{-10}$ | 1 | Log sampling |
| | L2 regularization penalty | $1e^{-10}$ | 1 | Log sampling |

## Notes

### Competing Interest Statement

AK has a patent for hollyTM (formerly Anxiety Meter) with royalties paid from Awake Labs. AK has received consulting fees from DNAStack and Shaftesbury. EA has received grants from Roche and Anavex, served as a consultant to Roche, Quadrant Therapeutics, Ono, and Impel Pharmaceuticals, has received in-kind support from AMO Pharma and CRA-Simons Foundation, received royalties from APPI and Springer, received an editorial honorarium from Wiley, and has a patent for hollyTM (formerly Anxiety Meter). The remaining authors have no potential conflicts of interest to report.

### Summary of Updates

The manuscript has been revised per comments from peer-reviews, primarily to the introduction and discussion.

## References

1 Kieling C, Buchweitz C, Caye A, Silvani J, Ameis SH, Brunoni AR et al. Worldwide prevalence and disability from mental disorders across childhood and adolescence: Evidence from the global burden of disease study. JAMA Psychiatry 2024; 81: 347–356.

2 Vergunst F, Commisso M, Geoffroy MC, Temcheff C, Poirier M, Park J et al. Association of childhood externalizing, internalizing, and comorbid symptoms with long-term economic and social outcomes. JAMA Netw Open 2023; 6: e2249568–e2249568.

3 Stevanovic D. Impact of emotional and behavioral symptoms on quality of life in children and adolescents. Quality of Life Research 2013; 22: 333–337.

4 Cannon TD, Sun F, Mcewen SJ, Papademetris X, He G, van Erp TGM et al. Reliability of neuroanatomical measurements in a multisite longitudinal study of youth at risk for psychosis. Hum Brain Mapp 2014; 35: 2424–2434.

5 Shen X, MacSweeney N, Chan SWY, Barbu MC, Adams MJ, Lawrie SM et al. Brain structural associations with depression in a large early adolescent sample (the ABCD study&#xae;). EClinicalMedicine 2021; 42. doi:10.1016/j.eclinm.2021.101204.

6 Whittle S, Vijayakumar N, Simmons JG, Allen NB. Internalizing and Externalizing Symptoms Are Associated With Different Trajectories of Cortical Development During Late Childhood. J Am Acad Child Adolesc Psychiatry 2020; 59: 177–185.

7 Schmaal L, Hibar DP, Sämann PG, Hall GB, Baune BT, Jahanshad N et al. Cortical abnormalities in adults and adolescents with major depression based on brain scans from 20 cohorts worldwide in the ENIGMA Major Depressive Disorder Working Group. Mol Psychiatry 2016; 22: 900–909.

8 Durham EL, Jeong HJ, Moore TM, Dupont RM, Cardenas-Iniguez C, Cui Z et al. Association of gray matter volumes with general and specific dimensions of psychopathology in children. Neuropsychopharmacology 2021; 46: 1333–1339.

9 Zhang Y, Xu B, Kim HH, Muetzel R, Delaney SW, Tiemeier H. Differences in cortical morphology and child internalizing or externalizing problems: Accounting for the co-occurrence. JCPP Advances 2022; 2: e12114.

10 Tejavibulya L, Rolison M, Gao S, Liang Q, Peterson H, Dadashkarimi J et al. Predicting the future of neuroimaging predictive models in mental health. Mol Psychiatry 2022; 27: 3129–3137.

11 Romer AL, Ren B, Pizzagalli DA. Brain structure relations with psychopathology trajectories in the ABCD study. J Am Acad Child Adolesc Psychiatry 2023; 62: 895–907.

12 Lipschutz R, Powers A, Minton ST, Stenson AF, Ely TD, Stevens JS et al. Smaller hippocampal volume is associated with anxiety symptoms in high-risk Black youth. Journal of Mood & Anxiety Disorders 2024; 7: 100065.

13 Andre QR, Geeraert BL, Lebel C. Brain structure and internalizing and externalizing behavior in typically developing children and adolescents. Brain Struct Funct 2020; 225: 1369–1378.

14 Muetzel RL, Blanken LME, Jan van der Ende, Hanan El Marroun, Shaw P, Sudre G et al. Tracking brain development and dimensional psychiatric symptoms in children: A longitudinal population-based neuroimaging study. American Journal of Psychiatry 2018; 175: 54–62.

15 Jarvers I, Kandsperger S, Schleicher D, Ando A, Resch F, Koenig J et al. The relationship between adolescents’ externalizing and internalizing symptoms and brain development over a period of three years. Neuroimage Clin 2022; 36: 103195.

16 Bos MGN, Peters S, Van De Kamp FC, Crone EA, Tamnes CK. Emerging depression in adolescence coincides with accelerated frontal cortical thinning. Journal of Child Psychology and Psychiatry 2018; 59: 994–1002.

17 Chung J, Teo J. Mental Health Prediction Using Machine Learning: Taxonomy, Applications, and Challenges. Applied Computational Intelligence and Soft Computing 2022; 2022: 9970363.

18 Rashid B, Calhoun V. Towards a brain-based predictome of mental illness. Hum Brain Mapp 2020; 41: 3468–3535.

19 Chen J, Tam A, Kebets V, Orban C, Ooi LQR, Asplund CL et al. Shared and unique brain network features predict cognitive, personality, and mental health scores in the ABCD study. Nature Communications 2022 13:1 2022; 13: 1–17.

20 Yeung HW, Stolicyn A, Buchanan CR, Tucker-Drob EM, Bastin ME, Luz S et al. Predicting sex, age, general cognition and mental health with machine learning on brain structural connectomes. Hum Brain Mapp 2023; 44: 1913–1933.

21 Lai MC, Kassee C, Besney R, Bonato S, Hull L, Mandy W et al. Prevalence of co-occurring mental health diagnoses in the autism population: A systematic review and meta-analysis. Lancet Psychiatry 2019; 6: 819–829.

22 Volkow ND, Koob GF, Croyle RT, Bianchi DW, Gordon JA, Koroshetz WJ et al. The conception of the ABCD study: From substance use to a broad NIH collaboration. Dev Cogn Neurosci 2018; 32: 4–7.

23 Alexander LM, Escalera J, Ai L, Andreotti C, Febre K, Mangone A et al. An open resource for transdiagnostic research in pediatric mental health and learning disorders. Sci Data 2017; 4: 170181.

24 Somerville LH, Bookheimer SY, Buckner RL, Burgess GC, Curtiss SW, Dapretto M et al. The Lifespan Human Connectome Project in Development: A large-scale study of brain connectivity development in 5–21 year olds. Neuroimage 2018; 183: 456–468.

25 Achenbach TM, Rescorla LA. Manual for the ASEBA school-age forms & profiles: An integrated system of multi-informant assessment. Aseba: Burlington, VT, VT, 2001.

26 Jacobson NS, Truax P. Clinical significance: A statistical approach to defining meaningful change in psychotherapy research. J Consult Clin Psychol 1991; 59: 12–19.

27 Fischl B. FreeSurfer. Neuroimage 2012; 62: 774–781.

28 Desikan RS, Se F, Fischl B, Quinn BT, Dickerson BC, Blacker D et al. An automated labeling system for subdividing the human cerebral cortex on MRI scans into gyral based regions of interest. Neuroimage 2006; 31: 968–980.

29 Rapee RM, Oar EL, Johnco CJ, Forbes MK, Fardouly J, Magson NR et al. Adolescent development and risk for the onset of social-emotional disorders: A review and conceptual model. Behaviour Research and Therapy 2019; 123: 103501.

30 Kushki A, Anagnostou E, Hammill C, Duez P, Brian J, Iaboni A et al. Examining overlap and homogeneity in ASD, ADHD, and OCD: A data-driven, diagnosis-agnostic approach. Transl Psychiatry 2019; 9. doi:10.1038/s41398-019-0631-2.

31 Vandewouw MM, Brian J, Crosbie J, Schachar RJ, Iaboni A, Georgiades S et al. Identifying replicable subgroups in neurodevelopmental conditions using resting-state functional magnetic resonance imaging data. JAMA Netw Open 2023; 6: e232066–e232066.

32 Morgado F, Vandewouw MM, Hammill C, Kelley E, Crosbie J, Schachar R et al. Behaviour-correlated profiles of cerebellar-cerebral functional connectivity observed in independent neurodevelopmental disorder cohorts. Translational Psychiatry 2024 14:1 2024; 14: 1–11.

33 Kushki A, Cardy RE, Panahandeh S, Malihi M, Hammill C, Brian J et al. Cross-diagnosis structural correlates of autistic-like social communication differences. Cerebral Cortex 2021; 31: 5067–5076.

34 Sadat-Nejad Y, Vandewouw MM, Cardy R, Lerch J, Taylor MJ, Iaboni A et al. Investigating heterogeneity across autism, ADHD, and typical development using measures of cortical thickness, surface area, cortical/subcortical volume, and structural covariance. Frontiers in Child and Adolescent Psychiatry 2023; 2: 1171337.

35 Lundberg SM, Allen PG, Lee S-I. A Unified Approach to Interpreting Model Predictions. Adv Neural Inf Process Syst 2017; 30.https://github.com/slundberg/shap (accessed 26 Feb2024).

36 Cardenas-Iniguez C, Gonzalez MR. Recommendations for the responsible use and communication of race and ethnicity in neuroimaging research. Nat Neurosci 2024; 27: 615–628.

37 Bahathiq RA, Banjar H, Bamaga AK, Jarraya SK. Machine learning for autism spectrum disorder diagnosis using structural magnetic resonance imaging: Promising but challenging. Front Neuroinform 2022; 16: 949926.

38 Lai MC. Mental health challenges faced by autistic people. Nature Human Behaviour 2023 7:10 2023; 7: 1620–1637.

39 Serra-Blasco M, Radua J, Soriano-Mas C, Gómez-Benlloch A, Porta-Casteràs D, Carulla-Roig M et al. Structural brain correlates in major depression, anxiety disorders and post-traumatic stress disorder: A voxel-based morphometry meta-analysis. Neurosci Biobehav Rev 2021; 129: 269–281.

40 Wu GR, Baeken C. Normative modeling analysis reveals corpus callosum volume changes in early and mid-to-late first episode major depression. J Affect Disord 2023; 340: 10–16.

41 Peng W, Jia Z, Huang X, Lui S, Kuang W, Sweeney JA et al. Brain structural abnormalities in emotional regulation and sensory processing regions associated with anxious depression. Prog Neuropsychopharmacol Biol Psychiatry 2019; 94. doi:10.1016/j.pnpbp.2019.109676.

42 Liang J, Yu Q, Liu Y, Qiu Y, Tang R, Yan L et al. Gray matter abnormalities in patients with major depressive disorder and social anxiety disorder: a voxel-based meta-analysis. Brain Imaging Behav 2023; 17: 749–763.

43 Liu X, Klugah-Brown B, Zhang R, Chen H, Zhang J, Becker B. Pathological fear, anxiety and negative affect exhibit distinct neurostructural signatures: evidence from psychiatric neuroimaging meta-analysis. Translational Psychiatry 2022 12:1 2022; 12: 1–19.

44 Zahn-Waxler C, Klimes-Dougan B, Slattery MJ. Internalizing problems of childhood and adolescence: Prospects, pitfalls, and progress in understanding the development of anxiety and depression. Dev Psychopathol 2000; 12: 443–466.

45 Mayes SD, Castagna PJ, Waschbusch DA. Sex differences in externalizing and internalizing symptoms in ADHD, autism, and general population samples. J Psychopathol Behav Assess 2020; 42: 519–526.

46 Horwitz E, Vos M, De Bildt A, Greaves-Lord K, Rommelse N, Schoevers R et al. Sex differences in the course of autistic and co-occurring psychopathological symptoms in adolescents with and without autism spectrum disorder. Autism 2023; 27: 1716–1729.

47 Herrmann L, Barkmann C, Bindt C, Fahrenkrug S, Breu F, Grebe J et al. Binary and non-binary fender identities, internalizing problems, and treatment wishes among adolescents referred to a gender identity clinic in Germany. Arch Sex Behav 2024; 53: 91–106.

48 Farhane-Medina NZ, Luque B, Tabernero C, Castillo-Mayén R. Factors associated with gender and sex differences in anxiety prevalence and comorbidity: A systematic review. Sci Prog 2022; 105. doi:10.1177/00368504221135469/ASSET/IMAGES/LARGE/10.1177_00368504221135469-FIG1.JPEG.

49 Garavan H, Bartsch H, Conway K, Decastro A, Goldstein RZ, Heeringa S et al. Recruiting the ABCD sample: Design considerations and procedures. Dev Cogn Neurosci 2018; 32: 16–22.

50 Chaarani B, Hahn S, Allgaier N, Adise S, Owens MM, Juliano AC et al. Baseline brain function in the preadolescents of the ABCD Study. Nat Neurosci 2021; 24: 1176–1186.

51 Lisdahl KM, Tapert S, Sher KJ, Gonzalez R, Nixon SJ, Feldstein Ewing SW et al. Substance use patterns in 9-10 year olds: Baseline findings from the adolescent brain cognitive development (ABCD) study. Drug Alcohol Depend 2021; 227: 108946.

52 Clark DB, Fisher CB, Bookheimer S, Brown SA, Evans JH, Hopfer C et al. Biomedical ethics and clinical oversight in multisite observational neuroimaging studies with children and adolescents: The ABCD experience. Dev Cogn Neurosci 2018; 32: 143–154.

53 Auchter AM, Hernandez Mejia M, Heyser CJ, Shilling PD, Jernigan TL, Brown SA et al. A description of the ABCD organizational structure and communication framework. Dev Cogn Neurosci 2018; 32: 8–15.

54 Diagnostic and statistical manual of mental disorders (DSM-5®). American Psychiatric Association, 2013.

55 Lord C, Rutter M, DiLavore PC, Risi S, Gotham K, Bishop S. Autism Diagnostic Observation Schedule, Second Edition (ADOS-2) Manual (Part I): Modules 1–4. Western Psychological Services, 2012.

56 Lord C, Rutter M, Le Couteur A. Autism Diagnostic Interview-Revised: A revised version of a diagnostic interview for caregivers of individuals with possible pervasive developmental disorders. J Autism Dev Disord 1994; 24: 659–685.

57 Kaufman J, Birmaher B, Brent D, Rao UMA, Flynn C, Moreci P et al. Schedule for Affective Disorders and Schizophrenia for School-Age Children-Present and Lifetime Version (K-SADS-PL): Initial reliability and validity data. J Am Acad Child Adolesc Psychiatry 1997; 36: 980–988.

58 Ickowicz A, Schachar RJ, Sugarman R, Chen SX, Millette C, Cook L. The Parent Interview for Child Symptoms: A situation-specific clinical research interview for attention-deficit hyperactivity and related disorders. The Canadian Journal of Psychiatry 2006; 51: 325–328.

59 Fischl B, Salat DH, Busa E, Albert M, Dieterich M, Haselgrove C et al. Whole brain segmentation: Automated labeling of neuroanatomical structures in the human brain. Neuron 2002; 33: 341–355.

60 Hagler DJ, Hatton SN, Cornejo MD, Makowski C, Fair DA, Dick AS et al. Image processing and analysis methods for the Adolescent Brain Cognitive Development Study. Neuroimage 2019; 202: 116091.

61 Glasser MF, Sotiropoulos SN, Wilson JA, Coalson TS, Fischl B, Andersson JL et al. The minimal preprocessing pipelines for the Human Connectome Project. Neuroimage 2013; 80: 105–124.

62 Dale AM, Fischl B, Sereno MI. Cortical Surface-Based Analysis: I. Segmentation and Surface Reconstruction. Neuroimage 1999; 9: 179–194.

63 Dale AM, Sereno MI. Improved localization of cortical activity by combining EEG and MEG with MRI cortical surface reconstruction: A linear approach. J Cogn Neurosci 1993; 5: 162–176.

64 Fischl B, Dale AM. Measuring the thickness of the human cerebral cortex from magnetic resonance images. Proc Natl Acad Sci U S A 2000; 97: 11050–11055.

65 Fischl B, Liu A, Dale AM. Automated manifold surgery: constructing geometrically accurate and topologically correct models of the human cerebral cortex. IEEE Medical Imaging 2001; 20: 70–80.

66 Fischl B, Salat DH, van der Kouwe AJW, Makris N, Ségonne F, Quinn BT et al. Sequence-independent segmentation of magnetic resonance images. Neuroimage 2004; 23: S69–S84.

67 Fischl B, Sereno MI, Dale AM. Cortical surface-based analysis: II. Inflation, flattening, and a surface-based coordinate system. Neuroimage 1999; 9: 195–207.

68 Fischl B, Sereno MI, Tootell RBH, Dale AM. High-resolution intersubject averaging and a coordinate system for the cortical surface. Hum Brain Mapp 1999; 8: 272–284.

69 Fischl B, van der Kouwe A, Destrieux C, Halgren E, Ségonne F, Salat DH et al. Automatically Parcellating the Human Cerebral Cortex. Cerebral Cortex 2004; 14: 11–22.

70 Han X, Jovicich J, Salat D, van der Kouwe A, Quinn B, Czanner S et al. Reliability of MRI-derived measurements of human cerebral cortical thickness: the effects of field strength, scanner upgrade and manufacturer. Neuroimage 2006; 32: 180–194.

71 Jovicich J, Czanner S, Greve D, Haley E, Van Der Kouwe A, Gollub R et al. Reliability in multi-site structural MRI studies: Effects of gradient non-linearity correction on phantom and human data. Neuroimage 2006; 30: 436–443.

72 Segonne F, Dale AM, Busa E, Glessner M, Salat D, Hahn HK et al. A hybrid approach to the skull stripping problem in MRI. Neuroimage 2004; 22: 1060–1075.

73 Reuter M, Rosas HD, Fischl B. Highly accurate inverse consistent registration: A robust approach. Neuroimage 2010; 53: 1181–1196.

74 Reuter M, Schmansky NJ, Rosas HD, Fischl B. Within-Subject Template Estimation for Unbiased Longitudinal Image Analysis. Neuroimage 2012; 61: 1402–1418.

75 Sled JG, Zijdenbos AP, Evans AC. A nonparametric method for automatic correction of intensity nonuniformity in MRI data. IEEE Trans Med Imaging 1998; 17: 87–97.

76 Segonne F, Pacheco J, Fischl B. Geometrically accurate topology-correction of cortical surfaces using nonseparating loops. IEEE Trans Med Imaging 2007; 26: 518–529.

77 Rosas HD, Liu AK, Hersch S, Glessner M, Ferrante RJ, Salat DH et al. Regional and progressive thinning of the cortical ribbon in Huntington’s disease. Neurology 2002; 58: 695–701.

78 Kuperberg GR, Broome M, McGuire PK, David AS, Eddy M, Ozawa F et al. Regionally localized thinning of the cerebral cortex in {S}chizophrenia. Arch Gen Psychiatry 2003; 60: 878–888.

79 Salat D, Buckner RL, Snyder AZ, Greve DN, Desikan RS, Busa E et al. Thinning of the cerebral cortex in aging. Cerebral Cortex 2004; 14: 721–730.

80 Fortin JP, Cullen N, Sheline YI, Taylor WD, Aselcioglu I, Cook PA et al. Harmonization of cortical thickness measurements across scanners and sites. Neuroimage 2018; 167: 104–120.

81 Bayer JMM, Thompson PM, Ching CRK, Liu M, Chen A, Panzenhagen AC et al. Site effects how-to and when: An overview of retrospective techniques to accommodate site effects in multi-site neuroimaging analyses. Front Neurol 2022; 13: 923988.

82 Ad-Dab’bagh Y, Einarson D, Lyttelton O, Muehlboeck J-S, Mok K, Ivanov O et al. The CIVET Image-Processing Environment: A Fully Automated Comprehensive Pipeline for Anatomical Neuroimaging Research. In: Proceedings of the 12th Annual Meeting of the Organization for Human Brain Mapping. NeuroImage: Florence, Italy, 2006.

83 Chollet F. Keras. GitHub repository. 2015.https://github.com/fchollet/keras (accessed 25 Feb2024).

84 Abadi M, Agarwal A, Barham P, Brevdo E, Chen Z, Citro C et al. TensorFlow: Large-scale machine learning on heterogeneous systems. 2015.https://www.tensorflow.org/.

85 Lemaître G, Nogueira F, Aridas CK. Imbalanced-learn: A python toolbox to tackle the curse of imbalanced datasets in machine learning. Journal of Machine Learning Research 2017; 18: 1–5.

